# Stress Granules Underlie Acute Myeloid Leukemia Stem Cell Survival and Stress Adaptation

**DOI:** 10.1101/2025.01.14.632811

**Authors:** Amanda Tajik, Emily Tsao, Soheil Jahangiri, Brendon Seale, Brian A. Yee, Jack T. Naritomi, Zaldy Balde, Severine Cathelin, Ava Keyvani Chahi, Lance Li, He Tian Chen, Nicholas Wong, Lina Liu, Pratik Joshi, Steven Moreira, Curtis W. McCloskey, Shahbaz Khan, Katherine L. Rothamel, Helena Boutzen, Suraj Bansal, Andy G.X. Zeng, Stefan Aigner, Yu Lu, John E. Dick, Thomas Kislinger, Rama Khokha, Mark D. Minden, Anne-Claude Gingras, Gene W. Yeo, Kristin J. Hope

## Abstract

The link between cancer maintenance and an ability to sustain continued growth through stresses conferred by the cancer state itself is growing. However, there are significant gaps in our understanding of how this stress is managed, particularly at the level of cancer initiating cells. Here, we identify proteins comprising the dynamic, stress-adaptive ribonucleoprotein complexes known as stress granules (SG) to be enriched among the factors essential for leukemic stem cell (LSC)-driven leukemic propagation. Focusing on core SG nucleator G3BP1, we dissect the role of SGs in human acute myeloid leukemia (AML), their targetability, and the mechanisms they govern to uncover a novel propensity for AML, and in particular LSC-enriched fractions, to prime the expression of SG components, form SGs with greater fidelity and to be reliant on their establishment and continued integrity for LSC maintenance. We further unveil the transcript and protein interactome of G3BP1 in the AML context and show that consolidated control of innate immune signaling, and apoptosis repression is executed through regional binding specificity of G3BP1 to highly structured 3’UTRs and cooperation with the RNA helicase UPF1 to mediate transcript decay in SGs. Altogether our findings advance novel fundamental principles of stress adaptation exploited in AML and LSCs that may extend to other cancers and uncover SGs as a novel axis for therapy development.

## INTRODUCTION

Acute myeloid leukemia (AML) is an aggressive blood cancer characterized by clonal expansion of immature myeloblasts which suppress healthy blood production. Initiation and maintenance of the disease is driven by leukemic stem cells (LSCs), the progeny of transformed hematopoietic stem or progenitor cells that have acquired superior self-renewal and perturbed differentiation^1^.

Sparing of LSCs by conventional treatment seeds disease resurgence in many cases, often in the form of highly aggressive and refractory relapses which underlie the poor 23% 5-year survival rate in AML^2–4^. Given the critical role of LSCs in both initiating and propagating leukemia, their effective targeting represents a key therapeutic goal, but one that is currently challenged by our limited understanding of the targetable molecular drivers of the LSC state and what underlies their invulnerability.

The initiation and progression of cancer is strongly connected to stress stimuli, exposing transformed cells to elevated levels of genotoxic, endoplasmic reticulum (ER), hypoxic, metabolic and oxidative stress^5,6^. Survival through stress exists in a balance, such that above a certain threshold of unresolved stress cells will initiate programmed cell death. However, the stress-preconditioned state of cancer cells appears to elevate their threshold for stress tolerance. Indeed, stress-adaptive responses, including the DNA damage response (DDR), unfolded protein response (UPR), autophagy and others, are co-opted in various cancers to promote survival and resistance to therapy^5,6^. In addition to these processes, which are enacted in response to specific stressors, the assembly of non-membranous organelles known as stress granules (SGs) has emerged as another axis that may present a critical avenue to mediate convergent stress adaptation and fitness-optimization in cancer^7^.

SGs are intracellular condensates comprised of ribonuclear proteins (RNP) that have assembled on mRNAs blocked from translation initiation, a process which can be triggered by a variety of physiological and pathological stressors, including the above-mentioned tumor-associated stimuli as well as externally imposed stress from UV radiation and chemotherapy^8^. SG formation, which thus far has primarily been shown in response to extrinsic insults, is initiated by core nucleating proteins, the foremost being G3BP stress granule assembly factor 1 (G3BP1), which is both indispensable for SG assembly and can activate SG nucleation when upregulated^9–11^. Although there remain many gaps in our understanding of how SGs function, mechanisms that have been reported include these G3BP1-dependent RNP complexes acting as scaffolds for selective sequestration or exclusion of signaling molecules to orchestrate dynamic signaling and survival throughout cellular stress^12,13^. SGs can also exert fine-tuned RNA-level control of gene expression by preferentially sequestering and stabilizing mRNAs coding for factors requiring rapid reactivation following stress cessation, while excluding those coding for chaperones and cell damage repair enzymes whose localization and continued translation at polysomes is essential for cellular integrity during stress^14,15^. By these means, SGs have been implicated as adaptive mechanisms in cancers including breast, colon, and pancreatic cancers, that contribute to disease progression and/or chemoresistance^15–17^. In AML however, a cancer known to exist in a highly inflammatory and thus inherently stressful state and driven by an LSC-population remarkably adept at evading therapy, a potential SG contribution has yet to be explored^18^. This paucity of understanding extends to the stem cell level where for LSCs and indeed cancer stem cells in general, a role for SGs has also been particularly overlooked.

Here we interrogate our recent high throughput in vivo assignment of RNA regulators as dependencies in AML and LSCs identifying that SG proteins are enriched among the factors essential for LSC-driven leukemic propagation^19^. By dissecting the role of G3BP1-dependent SGs in human AML and LSCs, their targetability, and the mechanisms they govern we uncover a novel propensity for AML, and in particular LSC fractions, to prime the expression of SG components, form SGs with greater fidelity and to be reliant on their formation and continued integrity for LSC maintenance. We further uncover an AML-specific G3BP1 SG proteome and transcriptome to show that orchestrated regulation of innate immune signaling, the fine balance of which is an emerging hallmark of myeloid neoplasms, and apoptosis repression is achieved through preferential regional binding of G3BP1 to highly structured 3’UTRs and cooperation with the RNA helicase UPF1 to enforce transcript decay in SGs. These insights unveil a novel paradigm of stress adaptation exploited in AML and LSCs and forward SGs as a promising therapeutic target.

## RESULTS

### SG proteins are elevated in human AML LSCs

In our previous study^19^ we performed CRISPR dropout screening in the LSC-driven RN2c leukemia (MLL-AF9^+^, NRasG12D^+^)^20^ to identify RNA binding proteins (RBPs) required for leukemic survival and propagation *in vivo*, including through secondary transplantation, a function uniquely achieved by self-renewal competent LSCs. Screen candidates were selected based on pronounced human LSC-specific enrichment with the goal of identifying the highest confidence therapeutic targets. Surprisingly, proteins of the Tier 1 consensus SG proteome (RNAgranuleDB v2.0)^21^ were significantly over-represented (**Fig. 1A**, hypergeometric test, p = 0.016) among the screen candidates that depleted upon secondary transplant (“secondary hits”), comprising 50% of those RBPs crucial for this LSC-specific function. For example, sgRNAs targeting the well-recognized G3BP-interacting SG protein, Caprin1, significantly depleted over serial transplantation (**Fig. 1B**). Prospective evidence of LSC impairment was apparent in primary grafts via significantly reduced expression of the LSC marker c-Kit in Caprin1 knockout cells compared to control (sgAno9) cells (**Fig. 1C and D**). We further analyzed the general essentiality of gene hits within the screen using the Core Essential Genes 2.0 (CEG2) set evaluated from genome scale CRISPR knockout screens across 17 cancer and immortalized cell lines^22^. Interestingly, none of the secondary transplantation dependencies, as compared to 9/20 of the genes that dropped out in primary transplantation, are recognized CEGs (**Supplementary Fig. S1A**). This result further highlights the secondary screen hits and SG RBPs as potentially therapeutically tractable AML-specific dependencies.

**Figure 1.**
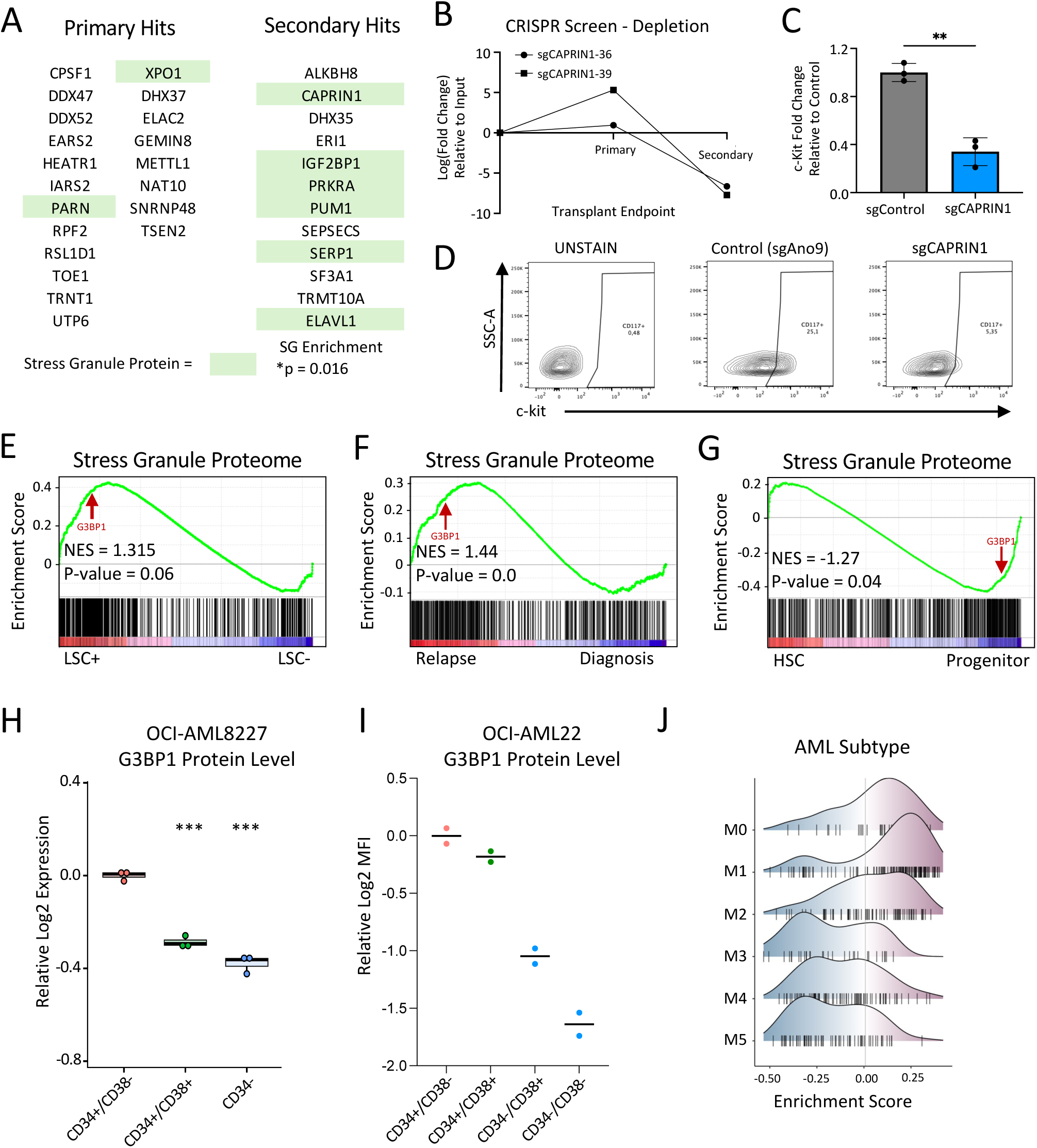
Stress Granule RBP Expression in Leukemia **A)** List of primary and secondary CRISPR dropout screen hits^19^. Stress granule proteins are highlighted in green. **B)** Log fold depletion of CAPRIN knockout guides in the primary and secondary transplants of the CRISPR dropout screen^19^. **C)** RN2c leukemia sgCAPRIN1-36 knockout results in a decrease of stem cell marker c-Kit upon primary transplantation endpoint. **D)** Representative Flow Cytometry plots depicting the loss of sgCAPRIN1 RN2c leukemia c-Kit expression upon primary transplantation. **E-G)** GSEA of the Jain et al^24^ SG proteome list using differential transcript profiles from **E)** functionally validated human LSC+ vs LSC-devoid fractions^23^, **F)** patient relapsed vs diagnostic samples^25^ and **G)** human bone marrow derived HSC vs progenitor fractions^26^. **H)** Proteomic expression of G3BP1 in OCI-AML-8227 stem (CD34^+^CD38^-^), progenitor (CD34^+^CD38^+^) and non-stem (CD34^-^) cells **I)** Median fluorescence intensity (MFI) from intracellular flow cytometric analysis of relative G3BP1 protein expression across the CD34/CD38 leukemic hierarchy. **J)** Density plots depicting Tier 1 SG proteome^21^ expression correlation analysis according to FAB blast morphology. ****P <.0001, ***P <.001, **P <.01, *P <.05. n.s. not significant.

Together our screen analyses provided the motivation to explore broadly the expression profiles of the SG proteome, and the core SG-nucleating factor G3BP1, in AML. Examining the expression profiles from 78 AML patients^23^ we uncovered a pronounced enrichment of the SG proteome^21,24^ in human AML LSCs compared to non-LSCs, as well as an enrichment within relapsed versus diagnosis AML samples^25^ (**Fig. 1E and F and Supplementary Fig. S1B and C**). Here, G3BP1 is present within the leading edge of these enrichments. In contrast, in healthy human bone marrow HSC versus progenitor expression profiles^26^ SG RBPs are downregulated in HSC populations (**Fig. 1G**). When considering G3BP1 RNA expression across HSCs and progenitor cells in human fetal liver, cord blood or bone marrow cells^26^, G3BP1 moreover demonstrates a pattern of increasing expression with decreased stemness of healthy tissue, aligned with the pattern observed for the overall SG proteome (**Supplementary Fig. S1D**). Furthermore, in patient AML models OCI-AML22 and OCI-AML-8227 each of which maintain a functional CD34^+^CD38^-^ LSC-driven leukemic hierarchy^27,28^, G3BP1 protein expression is greatest within the stem cell compartment compared to CD34^-^ bulk AML cells (**Fig. 1H and I and Supplementary Fig. S1E**).

To investigate SG associations with specific clinical subtypes further we looked at SG proteome expression across AML French America British (FAB) subtypes. SG proteome expression was found to be greatest among AMLs with more undifferentiated phenotypes, M0 and M1, consistent with an association with leukemic stem and progenitor cells (LSPCs) (**Fig. 1J**). Furthermore, there was no unique enrichment of SG proteins across AML subtypes classified by cytogenetic changes or common gene mutations, overall suggesting SGs are associated with LSCs regardless of AML background (**Supplementary Fig. S1F**). These expression profiles signal a heretofore unknown role of SGs in AML, in particular within the disease initiation and propagating fractions, providing the impetus to perform a systematic functional interrogation of AML and LSC SG dependencies in order to examine their potential as biomarkers and therapeutic targets.

### LSCs display heightened SG formation vs bulk AML

To determine the capacity of AML cells to induce SG formation we first validated immunofluorescence (IF) staining as well as live-cell imaging of transgenic EGFP-G3BP1 fusion constructs to visualize G3BP1^+^ puncta in THP-1 and OCI-AML22 AML cells subjected to prototypical SG induction by heat shock stress (**Fig. 2A and Supplementary Fig. S2A**). CellProfiler analysis of fixed and live-cell SGs showed a robust capacity for AML cells to induce SG formation, marked by G3BP1, following heat shock where both the number and size of SGs identified increase with stress (**Fig. 2B and C and Supplementary Fig. S2B and C**). In addition, using the live cell EGFP-G3BP1 SG reporter we observed a reduction in SGs following a return to control conditions showing the dynamic nature of SG formation and dissipation in AML (**Fig. 2D and E**).

**Figure 2.**
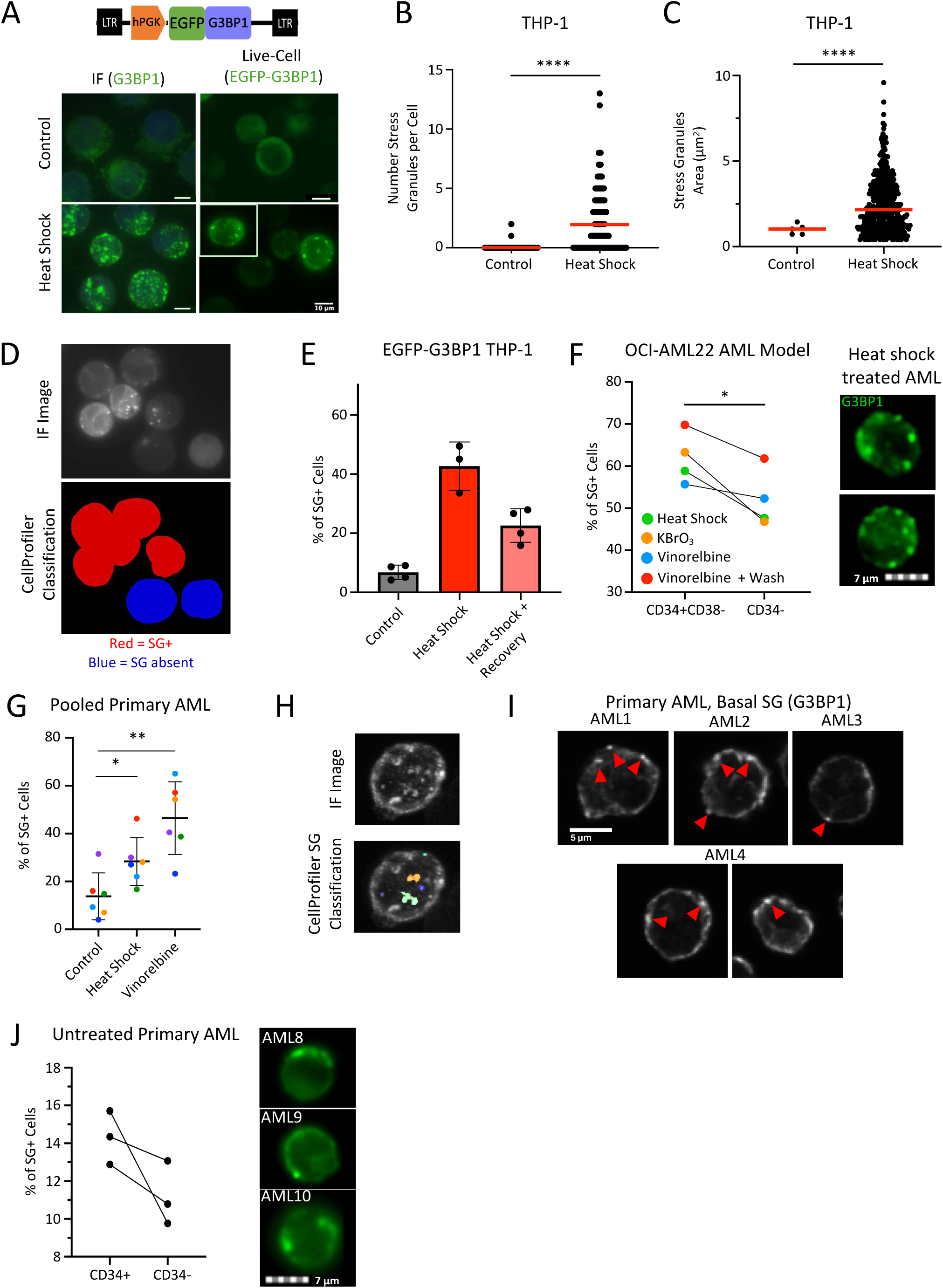
Stress granule dynamics in leukemia and normal HSPCs **A)** Schematic of lentiviral construct encoding EGFP-G3BP1 fusion (top) and IF and live-cell imaging of heat shocked THP-1 AML cells (bottom). IF images contain a nuclear stain (DAPI, blue) and G3BP1 (green) marks SG formation. SG formation in live-cell images was performed in stable cells expressing EGFP-G3BP1 fusion protein. **B-C)** Plots showing SG quantification per cell **B)** and average SG area **C)** of live-cell images of heat shocked (44 °C, 40 min) THP-1 cells. **D)** Cellprofiler output highlighting SG+ cells in red and SG-cells in blue. **E)** SG quantification using CellProfiler of stable EGFP-G3BP1 THP-1 cells following heat shock or recovery from heat shock. **F)** Representative Imagestream images of SG+ heat shocked OCI-AML22 cells (G3BP1 = green) (right) and plot showing percent of OCI-AML22 CD34^+^CD38^-^ LSC or CD34^-^ non-LSCs cells that were SG+ following heat shock, KBrO_3_ or vinorelbine treatments (left). **G)** Quantification of SG formation across patient AML specimens either untreated (basal level) or stimulated with heat shock or vinorelbine. Each symbol colour refers to a different biological sample (N=6). **H)** Representative image of primary AML heat shocked cells (top) and cellprofiler based identification of the presence of SGs in each cell (bottom). **I)** IF images of SGs in three different unstimulated primary AML samples. Red arrows point to G3BP1 puncta. **J)** Quantification of SGs in stem (CD34^+^) vs blast (CD34^-^) fractions of unstressed primary AML samples determined by Imagestream machine learning. Representative images depicting G3BP1 staining on the right. \*\*\*\**P* <.0001, \*\*\**P* <.001, \*\**P* <.01, \**P* <.05. n.s. not significant.

To probe for potential differences between leukemic stem vs blast populations in SG formation propensities we treated the functionally assessed LSC-driven OCI-AML22 AML patient model to SG-inducing heat shock, potassium bromate (oxidative stress), and vinorelbine (microtubule destabilizing stress) and used a custom ImageStream analysis pipeline on cells stained for surface CD34/CD38 and intracellular G3BP1 (**Supplementary Fig. S2D-F**). CD34^+^CD38^-^ OCI-AML22 LSCs demonstrated an increased formation of G3BP1^+^ SGs compared to CD34^-^ non-LSCs (**Fig. 2F**). Conversely, heat shocked healthy cord blood (CB) HSPCs demonstrate heightened SG formation in the more mature CD34^-^ fraction compared to CD34^+^CD38^-^ stem cells (**Supplementary Fig. S2G**).

Next, to translate these findings to patient samples we first validated by IF microscopy that primary AML cells robustly induce SG formation in response to heat shock and vinorelbine treatment (**Fig. 2G and H**). Surprisingly, we observed the presence of G3BP1^+^ puncta prior to the addition of extrinsic stress stimuli, suggesting SGs exist basally potentially to protect primary AML cells from chronic intrinsic stress (**Fig. 2I**). To dissect this at the level of LSPC vs blast fractions we used a machine learning algorithm trained to select for live and SG^+^ cells primarily based on nuclear morphology and G3BP1 contrast/homogeneity, respectively (**Supplementary Fig. S2H&I**). Quantification by this method showed a modest elevation of basal nucleated G3BP1^+^ puncta in untreated patient CD34^+^ LSPC fractions compared to CD34^-^ bulk fractions (**Fig. 2J**). Using the overall G3BP1 intensity levels in unstressed cells to estimate basal G3BP1 protein expression we found that in two out of three primary AML samples there were heightened G3BP1 levels in the CD34^+^ compartment (**Supplementary Fig. 2J**), whereas these appear reciprocally elevated in the CD34^-^ fractions of healthy CB samples (**Supplementary Fig. 2K**), supporting the concept that G3BP1 levels may play a role in enhancing the propensity towards SG formation. Altogether, our results indicate that AML LSPCs have a heightened propensity to nucleate SGs due to inherent oncogenic stress or extrinsic stress.

### AML blasts and LSCs are functionally dependent on SGs in vitro and in vivo

To investigate the functional effect of elevated G3BP1 levels in leukemia, we used low-level lentiviral overexpression of G3BP1. In MOLM-13 cells, increased G3BP1 expression led to improved competition in culture (**Supplementary Fig. 3A and B**), indicating a capacity for elevated G3BP1 to yield pro-growth effects on AML cells already highly proliferative. Next, to profile the functional dependency of leukemia cells on SGs, we used lentiviral delivery of shRNAs to knockdown G3BP1, a well-validated approach to diminish SG formation (**Fig. 3A and Supplementary Fig. S3C**)^17,29–31^. Knockdown of G3BP1 in THP-1 and MOLM-13 leukemia cells led to reduced growth in competitive cultures (**Fig. 3B and Supplementary Fig. S3D**) while flow cytometric analysis demonstrated increased differentiation (CD11b^+^) and apoptosis (**Fig. 3C and D**). Knockdown of another critical SG nucleator protein UBAP2L^10,29^ similarly resulted in decreased cell growth and elevated differentiation and apoptosis in THP-1 cells (**Supplementary Fig. S3E-H**). In addition, knockdown of G3BP1 or UBAP2L also yielded reduced growth of the OCI-AML22 patient model where we also observed reduced competition within the CD34^+^ LSC fraction demonstrating that the loss of requisite SG-nucleating proteins impairs AML propagation (**Supplementary Fig. S3I and J**).

**Figure 3.**
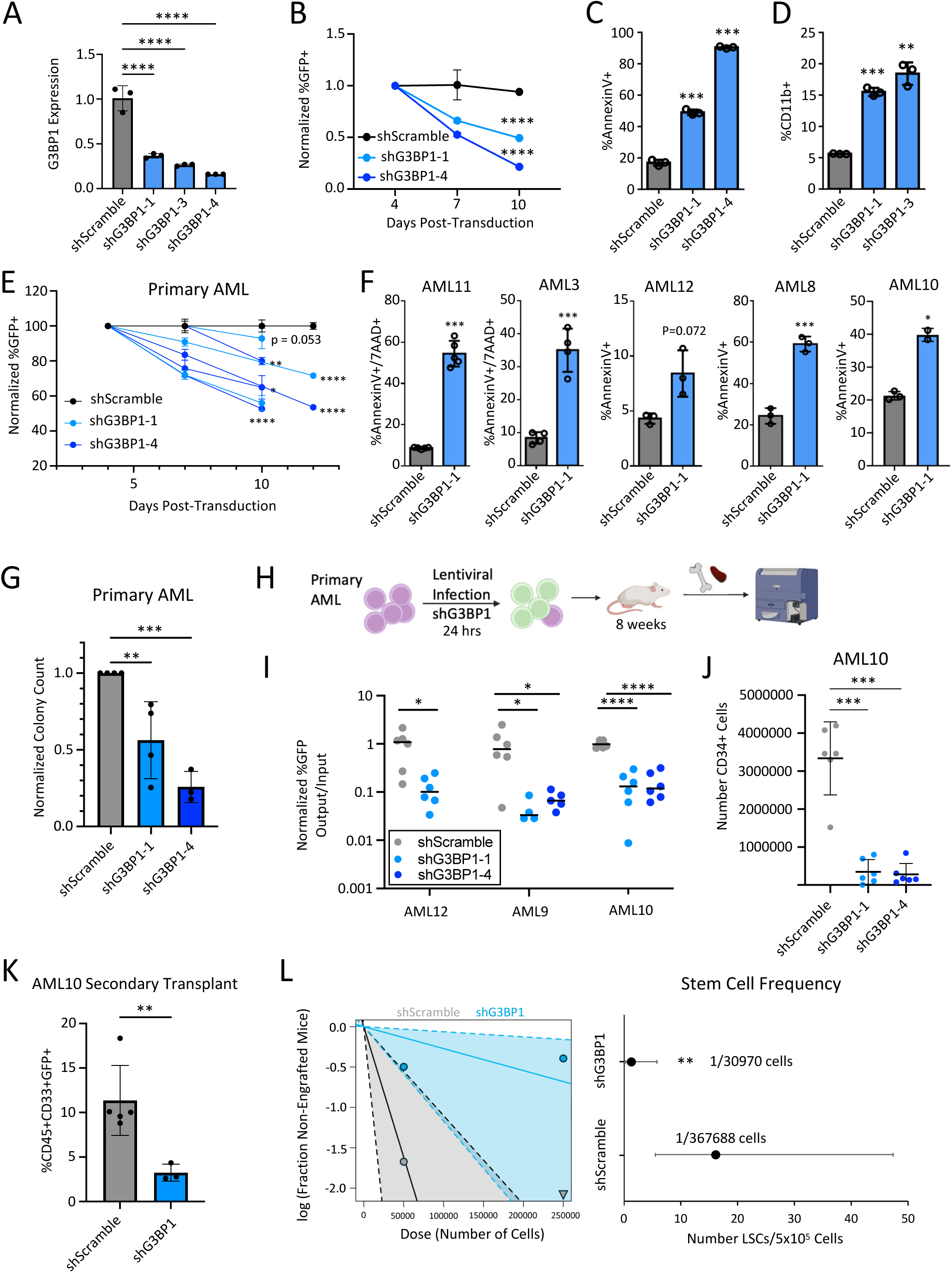
SG impairment via G3BP1 knockdown impairs LSC function in in vitro and in vivo assays **A)** qPCR validation of G3BP1 knockdown THP-1 cells normalized to GAPDH and shScramble (N=3). **B)** G3BP1 knockdown competitive cultures of MOLM-13 cells. **C-D)** G3BP1 knockdown THP-1 cell apoptosis **C)**, and differentiation marked by CD11b **D)**. **E)** Effect of G3BP1 knockdown on primary AML competitive growth in suspension culture (N=4 biological samples) **F)** and on apoptosis. **G)** Normalized colony forming unit assays of G3BP1 knockdown primary AML (N=4). **H)** In vivo xenotransplantation assay schematic. **I)** Engraftment of injected femur approximately 8 weeks after xenotransplantation of control (shScramble) or G3BP1 knockdown primary AML cells**. J)** CD34+ LSPCs in endpoint grafts formed by control vs G3BP1 knockdown primary AML. **K)** Engraftment of injected femurs of secondary mice by control and G3BP1 KD primary AML10 approximately 4 weeks after xenotransplantation and **L)** limiting dilution analysis of secondary recipients transplanted with BM of control vs G3BP1 knockdown primary AML. \*\*\*\**P* <.0001, \*\*\**P* <.001, \*\**P* <.01, \**P* <.05. n.s. not significant.

To translate these findings to patient samples, G3BP1-targeting shRNAs were lentivirally introduced into primary AML samples. Given that expression profiles prognosticate that SGs could be a pan-AML LSC dependency we evaluated AMLs across a diversity of cytogenetic backgrounds. We found that knockdown of G3BP1 decreased competition in culture, increased apoptosis and impaired AML progenitor output as measured by colony forming unit analysis in all specimens tested (**Fig. 3E-G and Supplementary Fig. S3K**). Depletion of UBAP2L similarly increased apoptosis of cultured primary AMLs (**Supplementary Fig. S3L**). To assess SG functions in LSC fractions, we performed gold-standard xenotransplantation of shG3BP1- transduced primary AML cells (**Fig. 3H**). At the transplant endpoint a significant loss of G3BP1 knocked down leukemia cells relative to controls was observed for all primary samples, including a highly aggressive relapsed AML sample (AML10) for which there was additionally a reduction in the number of CD34^+^ stem cell-enriched cells within the resulting graft (**Fig. 3I and J**). The essential requirement of SGs for LSC function was further validated with secondary transplant assays wherein CD45^+^CD33^+^GFP^+^ bone marrow grafts from primary control and G3BP1 knockdown mice were sorted and transplanted into secondary mice. Endpoint bone marrow analysis showed a significant impairment in secondary mouse engraftment capacity of G3BP1 knockdown cells (**Fig. 3K**). By carrying out these transplants using a high dose and low dose (250,000 and 50,000 cells respectively) we could apply limiting dilution analysis to determine a 12-fold reduction in peripherally engrafting LSCs following G3BP1 knockdown (**Fig. 3L**). These results suggest that the reduction of G3BP1 knockdown grafts in the primary transplant are due at least in part to decreased LSC self-renewal. Overall, this data demonstrates SG fidelity is an absolute requirement for leukemic cells including the disease-driving malignant stem cells.

### Small molecule inhibition of SG formation impairs growth and viability of human AML

To explore the potential efficacy of SG inhibition as an anti-AML therapy we tested small molecule inhibitors of G3BP1-mediated SG nucleation, resveratrol (RSVL) and G3Ib, for their therapeutic efficacy against AML. Both compounds have been reported to interact with G3BP1’s NTF2L domain, which is critical for G3BP dimerization and interaction with other key SG proteins, including CAPRIN1^32,33^.

RSVL (trans-3,5,4′-truhydroxystilbene) is a naturally occurring compound which through its interaction with G3BP1 can activate pro-apoptotic p53 signaling^34^. Others have also shown the interaction of RSVL with G3BP1 reduces the RNA-binding capacity of G3BP1^33^. However, as a direct demonstration of impaired SG assembly due to RSVL has not been shown, we used our stable THP-1 EGFP-G3BP1 fusion model and live-cell imaging to validate a robust decrease in SG formation in response to vinorelbine treatment when cells were cultured in the presence of RSVL compared to control DMSO (**Supplementary Fig. S4A-C**). AML cells treated with SG- inhibiting doses of RSVL demonstrated a dose-dependent decrease in in vitro cell growth, alongside increased apoptosis at early timepoints, compared to control DMSO treated cells (**Supplementary Fig S4D and E**). Treatment of the OCI-AML22 patient model further showed a reduction in the CD34^+^CD38^-^ LSC fraction upon RSVL treatment, mirroring the effects of G3BP1 knockdown (**Supplementary Fig. S4F**).

While RSVL can impede SG formation it does also have other described effects beyond SG regulation. In this respect G3Ib, a peptidomimetic resembling the FGDF motif of the viral nsP3 peptide represents a more specific inhibitor of SG formation through direct inhibition of G3BP1 NTF2L interactions^32^. Live-cell imaging of the THP-1 EGFP-G3BP1 fusion model indeed validated a robust decrease in SG formation in response to heat shock and vinorelbine when these AML cells were cultured in the presence of G3Ib compared to control DMSO (**Fig. 4A-C and Supplementary Fig. S4G**). THP-1 and OCI-AML22 cells treated with SG-inhibiting doses of G3Ib demonstrated a dose-dependent impairment of in vitro cell growth throughout culture associated with elevated apoptosis at early timepoints, compared to DMSO treated cells (**Fig. 4D and E**). In the OCI-AML22 AML model, after 7 days of culture there was an enhanced apoptotic effect in the CD34^+^ LSC fraction compared to CD34^-^ fraction, with a G3Ib dose-dependent depletion of LSPCs, further supporting a particularly high dependence of LSCs on SG fidelity (**Supplementary Fig. S4H**). Next, we examined the effect of 50 μM G3Ib on primary patient AML samples in vitro which resulted in increased apoptosis after 3 days in suspension culture (**Fig. 4F**) and impairment of colony forming capacity (**Fig. 4G and Supplementary Fig. S4I**). In contrast, parallel treatment of healthy CB specimens with 50 μM G3Ib had reduced impact on apoptotic status (**Fig 4H**) and no significant effect on total colony forming capacity or composition (**Fig. 4I**). Altogether these findings provide an important pre-clinical proof of concept that inhibition of SG nucleation via small molecules can selectively diminish the viability of AML and advances their potential for future anti-leukemic therapeutics development.

**Figure 4.**
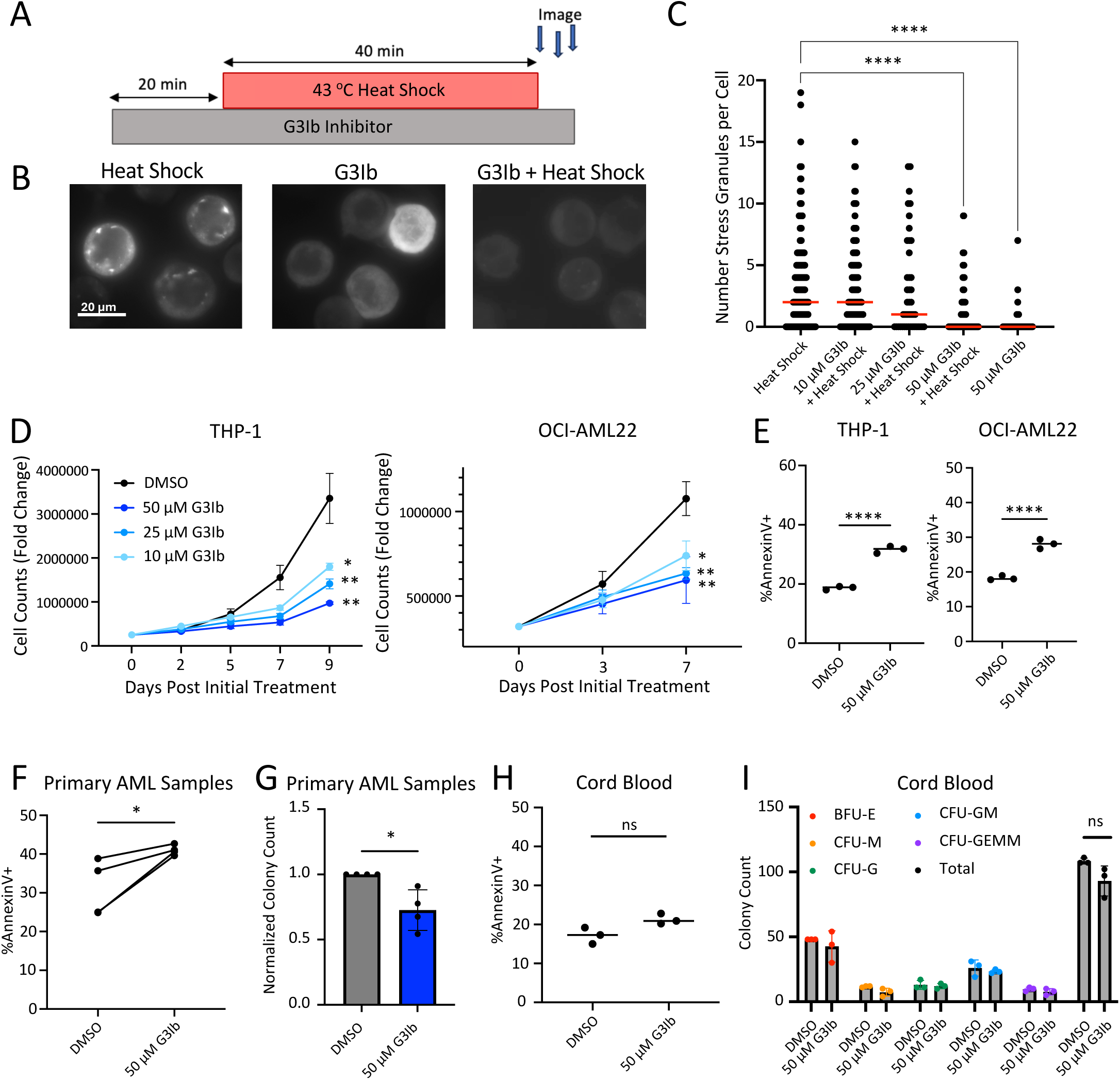
G3Ib chemical inhibition of SG formation impairs leukemic growth in culture **A)** THP-1 GFP-G3BP1 G3Ib inhibitor treatment schematic. **B)** Widefield microscope images of GFP-G3BP1 fusion THP-1 cells +/- G3Ib SG inhibitor and +/- heat shock (43 °C). **C)** Cellprofiler analysis of resulting #SG/cell. **D-E)** THP-1 and OCI-AML22 AML G3Ib treated or DMSO control treated fold change in culture **D)** and apoptosis after 2 and 3 days respectively. **E)**. **F)** Increased apoptosis in primary AML treated with 50 μM G3Ib (N=4 biological replicates) for 3 days. **G)** Impairment of AML colony forming capacity (N= 4 pooled analysis) when pre-treated with 50 μM G3Ib. **H)** Apoptosis staining in CB treated with G3Ib for 3 days (N=3). **I)** Colony forming capacity of 50 μM G3Ib pre-treated CB cells (N=3). \*\*\*\**P* <.0001, \*\*\**P* <.001, \*\**P* <.01, \**P* <.05. n.s. not significant.

### SGs control multiple critical pathways in AML LSCs at the RNA level

While there are many canonical players within SGs it is also clear that cell-and stress-context specific factors can be critical contributors to unique disease pathologies^35,36^. This framework of SG-associated proteins and RNAs are uncharacterized in AML. Thus, in order to gain insights into the function of G3BP1 and SGs in governing gene expression in AML and LSCs we used a combination of gene expression and RNA-binding mapping approaches. First, we performed paired transcriptomics (RNA-seq) and proteomics on G3BP1 knockdown THP-1 cells, as well as RNA-seq on the CD34^+^ fraction of five primary AML samples following G3BP1 knockdown. Upon G3BP1 loss, RNA-seq and proteomics results were positively correlated demonstrating G3BP1 stabilized transcripts are, as expected, upregulated at the protein level and conversely, transcripts destabilized by G3BP1 have commensurately reduced protein levels (**Fig. 5A and Supplementary Fig. S5A**). Pathway analysis of the transcriptional datasets showed downregulation of MYC targets, Cell Cycle, DNA Repair, Translation and mRNA Splicing upon G3BP1 reduction in THP-1 and primary AML (**Fig. 5B and Supplementary Fig. S5B**). Consistent with expression changes of these pathways, Ki67 staining showed G0 stalling of cell cycle and reduced global protein synthesis as measured by OP-Puro incorporation levels in G3BP1 knockdown THP-1 cells (**Supplementary Fig. 5C and D**). Conversely, signatures upregulated with G3BP1 knockdown include genes with lower expression in HSCs, and higher expression in myeloid cell development. Moreover, GO terms corresponding to Apoptosis and a collection of immune system related signatures including Innate Immune System, Signaling by Interleukins, Inflammatory Response, Interferon Gamma Response, and Interferon Alpha Response were also enriched in G3BP1 knockdown gene expression profiles (**Fig. 5B and C**). Importantly, G3BP1 reduction also resulted in a loss of the primitive LSC transcriptional signature^23^ in primary AML (**Fig. 5D**), which was also seen at the proteome level in THP-1 AML cells (**Fig. 5E**) and consistent with the outcomes of our functional assays indicating a decrease in stem cell properties. By analyzing G3BP1 gene expression correlation data from a consortium of RNA-seq of patient AML samples (CBioPortal – OHSU, 2022), we observed similar enrichments as in our G3BP1 knockdown results wherein low G3BP1 expression clinically correlates with heightened expression of apoptosis and inflammatory response genes (**Supplementary Fig. S5E**), further supporting the connection of these mechanisms in the greater patient population^37–40^. Spearman’s Correlation analysis comparing G3BP1 expression to genes in the apoptosis or inflammatory gene sets across a variety of cancers further shows this G3BP1-apoptosis/inflammation anticorrelation to be nearly unique and most prominent in AML (**Supplementary Fig S5F and G**).

**Figure 5.**
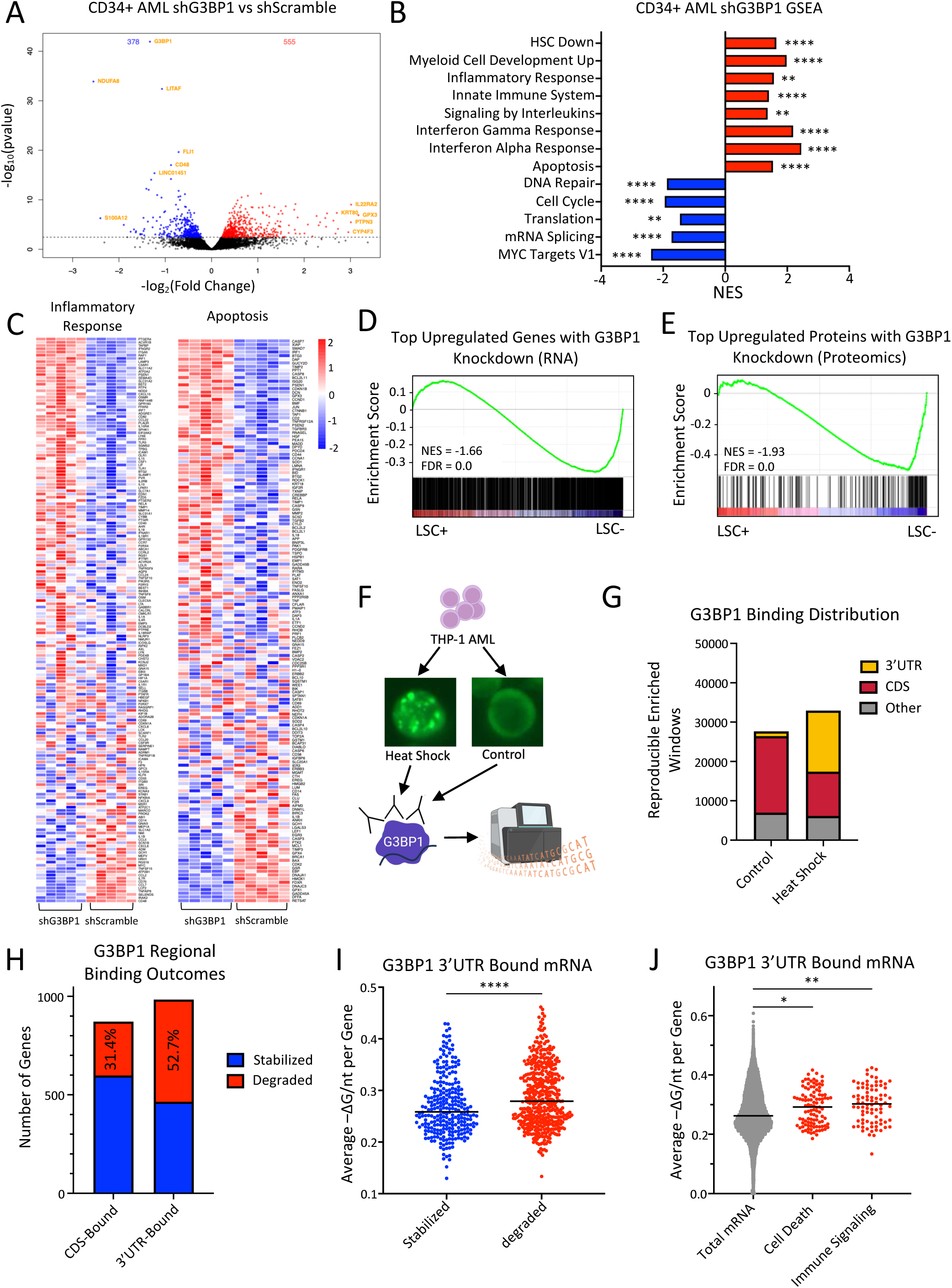
Multi-omic analysis of G3BP1 SG post-transcriptional regulation. **A)** Volcano plot of gene expression changes in the CD34^+^ fraction of 5 primary AML samples upon G3BP1 knockdown and **B)** barplot showing notable positively and negatively enriched pathways. **C)** Heat maps showing gene expression changes in G3BP1 knockdown primary AML cells within apoptosis and inflammatory genesets. **D)** GSEA of the top upregulated genes in G3BP1 knockdown primary AML transcriptome compared against transcript profiles from patient LSC+ vs LSC-fractions. **E)** GSEA of the top upregulated proteins in G3BP1 knockdown THP-1 proteome compared against transcript profiles from patient LSC+ vs LSC- fractions. **F)** Schematic of THP-1 treatment and eCLIP-seq workflow. **G)** Comparison of regional binding of G3BP1 from eCLIP-seq of untreated vs heat shocked THP-1 **H)** Barplot showing the fraction of genes bound at the CDS and 3’UTR in the THP-1 heat shock condition associated with RNA decay or stabilization from G3BP1 KD RNA-seq analysis. **I)** Minimal thermodynamic free energy of 3’UTRs for transcripts bound by G3BP1 in 3’UTRs and either stabilized or degraded. G3BP1 eCLIP 3’UTR bound mRNA associated with RNA decay have greater overall 3’UTR structure compared to stabilized transcripts. **J)** Minimal thermodynamic free energy of 3’UTRs for transcripts involved in cell death or immune signaling gene sets that are bound in 3’UTRs and degraded by G3BP1. \*\*\*\**P* <.0001, \*\*\**P* <.001, \*\**P* <.01, \**P* <.05. n.s. not significant.

To understand how these programs are directly intersected with G3BP1 and SGs, we mapped G3BP1 RNA binding profiles using enhanced cross-linking immunoprecipitation sequencing (eCLIP-seq) on control vs stressed SG^high^ (heat shocked) THP-1 AML cells (**Fig. 5F**). As well as capturing transcripts bound, eCLIP-seq further identifies the region of transcripts bound by G3BP1. Using this approach, we found that the majority of G3BP1 binding occurred in mRNA CDS and 3’UTRs, and a striking 10.7-fold increase in 3’UTR association took place in the SG^high^ condition compared to control, suggesting a stress-mediated regional bias in G3BP1 RNA associations within SGs (**Fig. 5G**). This switch in G3BP1 mRNA regional binding from the CDS to 3’UTR has been reported in other types of stressed cells, however it appears in the AML context to increase to a surprisingly greater extent. For example, G3BP1 3’UTR binding increased only 2.4-fold in puromycin stressed pluripotent stem cell-derived motor neurons^36^. Additionally, previous reports from other cell contexts have shown that transcripts preferentially associated within SGs contain reduced total transcript GC composition and 3’UTRs with greater length, patterns that we also observed in our 3’UTR control vs. SG^high^ G3BP1-binding sets. (**Supplementary Fig. S6A and B**)^41,42^.

By integrating RNA-seq and eCLIP-seq outputs to examine transcript fates as a function of regional binding of G3BP1, we found that the majority of CDS-bound transcripts were downregulated upon G3BP1 depletion (**Fig. 5H**), indicating that these transcripts are likely targets of G3BP1-stabilization and regulated in accordance with the historical understanding of SGs and G3BP1 as primarily functioning to stabilize RNA. Intriguingly however, greater than half of the transcripts bound at the 3’UTR were upregulated in G3BP1 knockdown cells. In particular, we found that transcripts with longer 3’UTRs and higher GC content 3’UTRs were highly associated with upregulation in G3BP1 knocked down cells, indicating they are preferential candidate targets of G3BP1-mediated destabilization and repression (**Supplementary Fig. S6A and B**). Looking beyond length and GC parameters we explored whether the attribute of RNA structure might influence G3BP1 RNA binding and resultant outcomes. Intriguingly, the net effect of significantly increased 3’UTR RNA structure estimated from minimal thermodynamic free energy^43^, which was not a strong predictor of overall G3BP1-binding, was associated with target repression, pointing to a pattern of structure-associated decay of this particular set of G3BP1 targets in AML (**Fig. 5I & Supplementary Fig. S6C)**. This apparent structure-associated decay is seen in both the SG^high^ and control conditions, although the number of targets regulated in this manner increases following large-scale SG assembly. Exemplifying the binding-stabilization mode, MYC mRNA was bound by G3BP1 at the 3’UTR and CDS but has a 3’UTR structure below average, and consistent with the parameters of this model *MYC,* and MYC targets, were downregulated in G3BP1-depleted transcriptomes indicating a reliance on G3BP1 for *MYC* stabilization. In contrast, transcripts belonging to key upregulated Cell Death and Immune Signaling signatures upon G3BP1 knockdown have relatively highly structured 3’UTRs (**Fig. 5J**), consistent with these classes of transcripts being specifically sensitive to context-dependent SG-orchestrated structure mediated decay.

We next tested if these patterns of regional binding regulation could be seen in patient AML by performing eCLIP-seq on primary samples either untreated or following additional heat shock stress. First, we observed increased 3’UTR and decreased CDS binding under heightened stress conditions, mirroring the directionality of stress-induced shifts in binding seen in our THP-1 results (**Supplementary Fig. S7A)**. Importantly, when integrating gene expression outcomes we again found that association with 3’UTRs is more predictive of transcript repression compared to CDS-binding **(Supplementary Fig. S7B)**, and that features of bound 3’UTR length, GC content and overall structure are associated with G3BP1-mediated transcript degradation **(Supplementary Fig. S7C-F)**, reflecting the shared use of mechanisms uncovered in THP-1 cells by primary AML cells.

### Uncovering an AML-specific SG proteome

Given the central identity of SGs as a nucleating hub of protein interactions and evidence supporting context specificities of these interactions, we sought to uncover the specific protein interaction network nucleated by G3BP1 in AML using BioID. We generated lentiviral fusions of G3BP1 with the abortive biotin ligase miniTurbo to facilitate rapid, high accuracy biotinylation of proteins in close proximity to G3BP1 and their recovery and identification through subsequent streptavidin pulldown and mass spectrometry. Further stringency was provided in our experiment by the insertion of a self-cleaving P2A peptide between G3BP1 and the miniTurbo protein to facilitate their independent co-overexpression as a control (**Fig. 6A and B and Supplementary Fig. S8A**). The SG proteome was assessed in THP-1 and MOLM-13 leukemia cells, the CD34/CD38 stem, progenitor and bulk fractions of the OCI-AML22 AML model as well as across CD34^+^ and bulk patient AML samples. We first identified a stringent core set of G3BP1 interactors as those identified in at least six (out of eight total) AML samples. This yielded a network of 21 proteins which, as expected, was primarily composed of Tier 1 SG proteins such as G3BP2, UBAP2L and CAPRIN1 with a highly connected clustering coefficient in STRING (0.697, p<1.0e-16) (**Fig. 6C and D**). In accordance, GO:CC terms Cytoplasmic Ribonucleoprotein Granule and Cytoplasmic Stress Granule were the topmost enriched terms (**Supplementary Fig. S8B**). Next, to explore the possibility of an AML-unique SG protein network we examined the profile of interactors detected in a minimum of three AML samples. This uncovered >800 proteins and although the minority (<25%) of factors here are considered part of the canonical SG proteome^21^ these proteins together still had a significant STRING clustering coefficient, importantly indicating non-random network composition (0.376, p<1.0e-16) (**Fig. 6E and Supplementary Fig. S8C**). While we would expect variation between our G3BP1 specific AML interactome and the broader SG proteome, this result highlights a significant group of potentially AML specific SG proteins. Pathway overrepresentation analysis again showed an enrichment of GO:CC term Cytoplasmic Stress Granule (FDR= 3.20e-12), as well as a number of other key pathways in AML regulation including mRNA Splicing, Cell Cycle, Apoptosis, and Innate Immunity pathways (**Fig. 6F and Supplementary Fig. S8D**). This finding complements the results of our shG3BP1 RNA-sequencing and eCLIP-seq in support of consolidated regulation at both the RNA and protein level of these key processes in AML via SGs.

**Figure 6.**
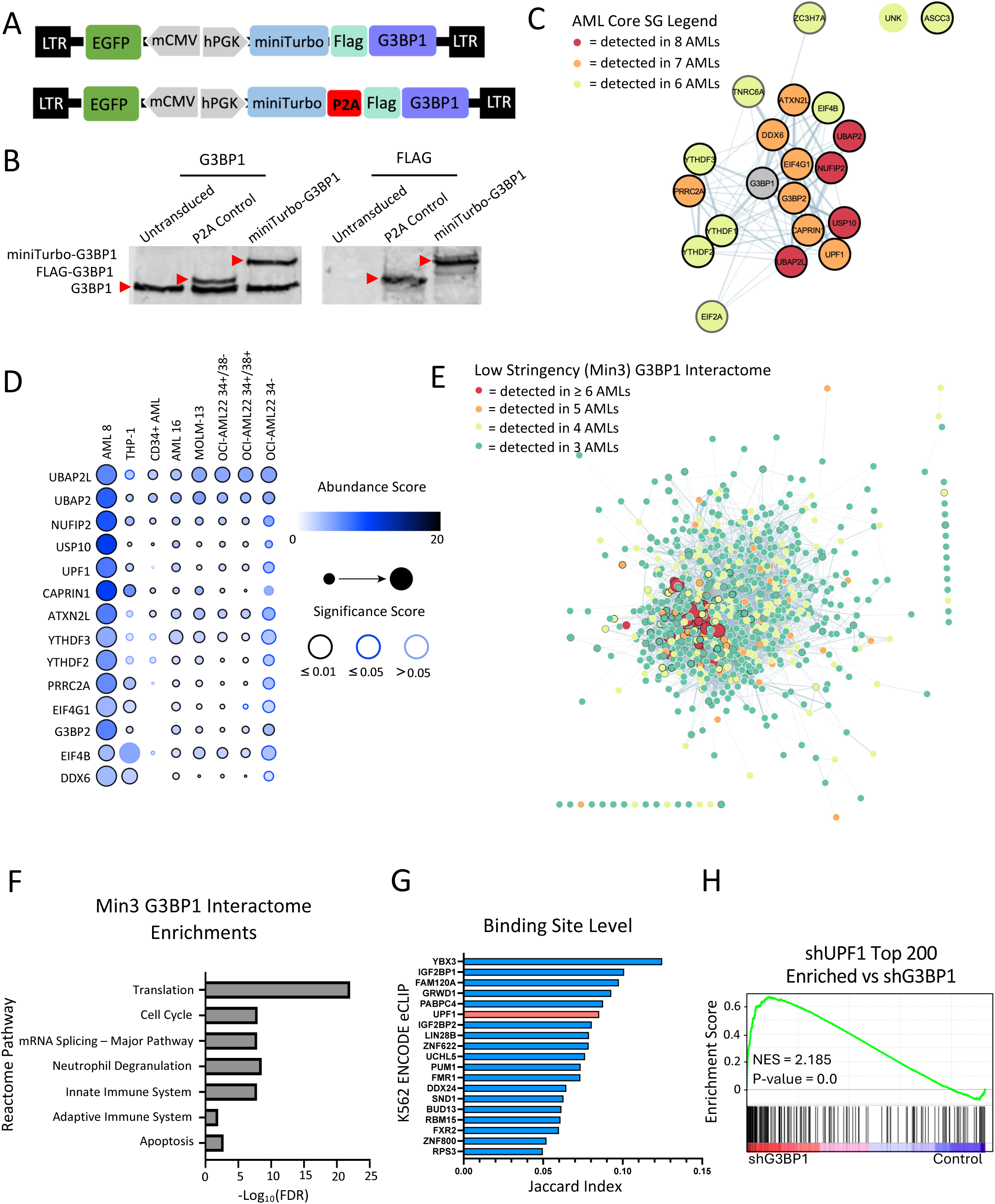
G3BP1 miniTurbo and the identification of the AML SG proteome **A)** Schematics of BioID lentivectors. **B)** Western blot validation of miniTurbo-3xFLAG-G3BP1 and P2A control fusions. **C)** STRING analysis of the core G3BP1+ SG proteome identified in a minimum of 6 (Min 6) AML G3BP1 BioID assays. **D)** Dot plot of core AML G3BP1 BioID hits. AML 8 and THP-1 cells were analyzed by DDA spectral counting and SaintExpress analysis abundance score provided is the fold change relative to control and the significance score provided in the Bayesian False Discovery Rate (BFDR). The remaining samples were analyzed via DIA peak area intensity and abundance score provided is the Log2(Fold Change) and q value significance score. **E)** STRING analysis of the G3BP1+ SG proteome identified in a minimum of 3 (Min 3) AML G3BP1 BioID assays. **F)** Significantly overrepresented Reactome pathways determined in STRING for the AML Min 3 SG proteome. **G)** Top K562 ENCODE^44^ RBPs holding a similar overlap in CLIP transcript binding site profiles to THP-1 heat shocked cells. **H)** GSEA plot showing enrichment of the top 200 upregulated genes upon UPF1 KD in the G3BP1 knockdown transcriptome.

To explore coordinated regulation between G3BP1 and its interactors in the AML context, we first performed comparative analysis between our SG^high^ AML G3BP1 eCLIP or post-G3BP1 knockdown RNA profiles and the publicly available ENCODE consortium’s RBP eCLIP and RBP knockdown RNA-seq datasets for >100 RBPs^44,45^. First, we found that the median global eCLIP overlap scores (Jaccard Index) increases when all RBPs were filtered for those identified in our Min3 and Min4 AML SG proteomes (**Supplementary Fig. S8E, columns 1-3**). The binding overlap scores further improve when considering RBPs showing similar differential gene expression profiles consistent with RBP-dependent degradation upon knockdown as compared to G3BP1 knockdown (**Supplementary Fig. S8E, columns 4-5**). Moreover, these indices of co-regulation displayed better performance in the K562 leukemic cell line compared to the HEPG2 liver cancer cell line, supporting the identification of a leukemia-selective SG network. This comparative analysis provides support for general coordination of gene expression regulation by the interacting RBPs identified within the AML SG. To assign the key players potentially synergizing with G3BP1 in AML SGs we next queried individual binding site and transcript level overlap scores (**Fig. 6G and Supplementary Fig. S8F**). This analysis revealed UPF1, which is part of the AML core SG network, to be a factor with highly similar transcript binding and gene regulatory profiles. We were particularly interested in UPF1 as it is an RNA helicase that has recently been shown to coordinate with G3BP1 to broadly facilitate decay of highly structured 3’UTR containing transcripts in an apparently SG independent manner in solid-tumor derived immortalized cell lines^43^. Importantly, comparing to our G3BP1-depleted primary AML RNA-seq, we indeed found a correlation with the top 200 upregulated genes in UPF1 knocked down K562s, highlighting the intriguing possibility that a G3BP1-UPF1 structure-mediated decay mechanism within SGs could be a vulnerability in patient AMLs (**Fig. 6H**).

### G3BP1 coordinates with UPF1 for the structure-mediated decay of pro-apoptotic transcripts in AML stress granules

Together our AML SG proteome and transcriptome coupled with gene expression outcomes herald a pro-survival mechanism whereby G3BP1^+^ SGs consolidate the downregulation of apoptotic and inflammatory transcripts via association with highly structured 3’UTRs and coordination with UPF1 to direct transcript decay. To evaluate this functionally, we first performed UPF1 knockdown competition assays where we observed that UPF1 depletion phenocopied G3BP1 knockdown displaying significantly decreased competition in culture (**Fig. 7A and B and Supplementary Fig. S9A**). To address whether UPF1 and G3BP1 were critical to the degradation of highly structured 3’UTR containing transcripts identified in our G3BP1 eCLIP assays, we performed Actinomycin D RNA decay assays in THP-1 cells with either G3BP1 or UPF1 knockdown. Selected target candidates were those bound by G3BP1 at the 3’UTR in the SG^high^ and/or control settings, had highly structured 3’UTRs (>0.27-ΔG/nt^43^) and were elevated upon G3BP1 knockdown. Focusing on the coordinated regulation of apoptosis regulators we evaluated the degradation dynamics of BCL2L11, DAP, and APAF1. BCL2L11 is a pro-apoptotic member of the BCL-2 protein family implicated in AML venetoclax resistance mechanisms^46^. DAP is an emerging positive regulator of apoptosis though currently little is known of its role in AML^47^. Finally, APAF1 is a component of the apoptosome that assembles following cytochrome c release into the cytoplasm and has been shown to be under negative regulation via DNA methylation in AML^48,49^. Importantly, upon G3BP1 or UPF1 knockdown, all three candidates showed decreased decay indicating that both G3BP1 and UPF1 are involved in the degradation of these transcripts (**Fig. 7C and D and Supplementary Fig. S9B**). Furthermore, we did not see a reduction in decay rates in control RNA candidates bound by G3BP1 with low 3’UTR structures (<0.27 -ΔG/nt) (**Supplementary Fig. S9C**).

**Figure 7.**
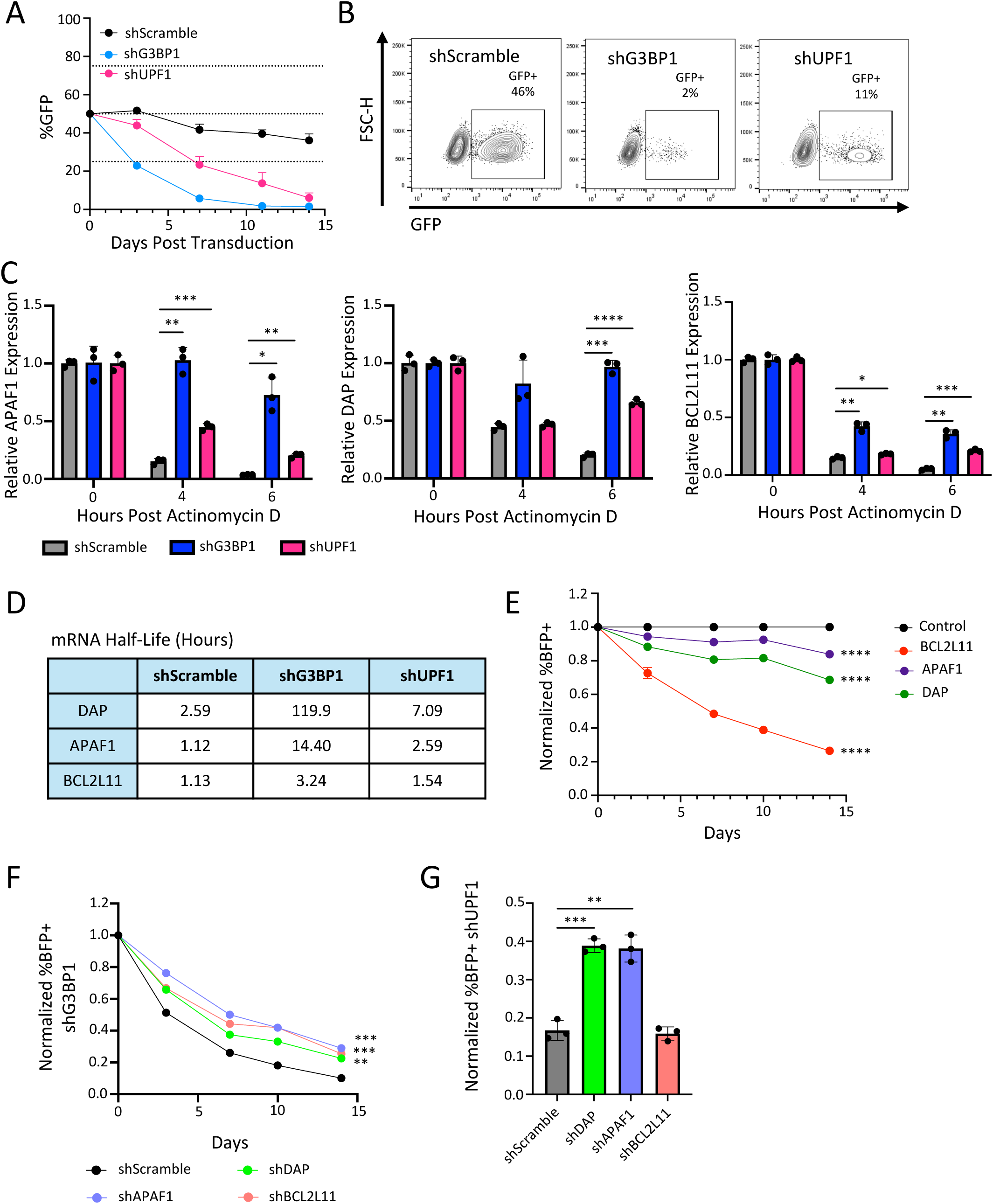
G3BP and UPF1 Coordinate in the structure-mediated decay of pro-apoptotic transcripts **A)** shG3BP1-4 and shUPF1-2 competitive cultures in THP-1 AML cells. **B)** Representative flow plots showing the loss of G3BP1 and UPF1 knockdown THP-1 cells versus shScramble cells after 11 days of competition. **C)** APAF1 (left), DAP (middle), and BCL2L11 (right) qPCR expression in shScramble, shG3BP1-4, or shUPF1-2 knockdown THP-1 cells 0, 3, and 6 hours post Actinomycin D treatment. **D)** Table of mRNA half-lives upon shScramble, shG3BP1-4, or shUPF1-2 knockdown. **E)** THP-1 competitive assays upon control overexpression relative to BCL2L11, APAF1, or DAP overexpression. **F)** THP-1 shG3BP1-4 competitive growth in culture when co-expressing shRNAs against DAP, BCL2L11, APAF1, or Scramble. **G)** Representation of BFP+ shUPF1-3 THP-1 cells in culture 3 days post double transduction to co-express shRNAs against DAP, BCL2L11, APAF1, or Scramble. \*\*\*\**P* <.0001, \*\*\**P* <.001, \*\**P* <.01, \**P* <.05. n.s. not significant.

To investigate whether the upregulation of these pro-apoptotic genes targeted for decay by G3BP1/UPF1 contributes to the AML-inhibitory phenotype of G3BP1 knockdown, we performed lentiviral overexpression assays. Indeed, we found that heightened levels of BCL2L11, APAF1, and DAP in THP-1 AML cells each resulted in significantly decreased competition of transduced cells in culture compared to control cells (**Fig. 7E**). To reciprocally determine if the repression of these apoptotic targets could rescue the G3BP1/UPF1 knockdown phenotype we utilized a candidate knockdown strategy to simulate the targeted decay expected to be enacted by G3BP1/UPF1. Knocking down each of these candidates allowed for partial rescue of both G3BP1 and UPF1 knockdown cells in competitive cultures, showing that the decay of these apoptotic transcripts by G3BP1/UPF1 is key to the maintenance of leukemia cells (**Fig. 7F and G and Supplementary Fig. S9D-G**). Overall, our AML G3BP1+ SG data demonstrates that SGs are essential to the survival and intrinsic stress adaptation response in AML through the structure-mediated suppression of pro-apoptotic transcripts.

## DISCUSSION

The link between cancer maintenance and an ability to sustain continued growth through stresses conferred by the cancer state itself is growing however there are significant gaps in our understanding of how this stress is managed in cancers whose origins position them to be stress prone^5,6^. Similar questions exist with respect to cancer initiating cells whose imperviousness to stress is paramount for survival of the entire cancer clone and are thus arguably the most requiring of robust stress-resistance mechanisms. Despite this, our understanding of the physiological relevance of SGs in these driver cells is lacking. Here we have discovered a unique transcriptional upregulation of the SG machinery as a whole, including the core essential nucleator G3BP1, in LSC-enriched fractions of AML which imparts both a heightened propensity towards mounting a protective SG response following insult as well as basal nucleation of SGs in patient samples. We show that this physiological elevation in SG levels is essential not just for the continued survival of leukemic blasts and progenitors but is fundamentally critical for integrity of the LSC pool at steady state. Our work demonstrates that G3BP1-SGs govern AML cell fate through a unique program of regional binding and structure-mediated decay to shield against overactivated apoptotic and innate immune signaling and are thereby functionally indispensable to mitigate the intrinsic stressors faced by AML cells and LSCs in particular. That our findings extend across AML agnostic to underlying mutational status speaks to our identification of SG addiction as a convergent oncogenic mechanism in AML. From a translational perspective, using a highly selective test compound to inhibit G3BP1-mediated SG nucleation, we also provide important proof of principle that SG targeting could present an efficacious anti-AML strategy.

Our finding of SG dependency across AML irrespective of genetic alterations highlights the need for robust stress resistance mechanisms in the development and maintenance of AML specifically. While other cancer cell types have also been shown to rely on SGs, it might be anticipated that the dependency of a given cancer on this axis will vary as a function of the extent of the underlying stress these cells are operating under chronically. Indeed, in pancreatic cancer, the added stress imposed by pre-existing obesity has been shown to confer an increased requirement for SGs in response to acute stressors^17^. In AML, inflammation is an inherent stressor by virtue of its myeloid lineage of origin and the elevated signaling that is enforced through further wiring of the leukemic epigenome and transcriptome^18^. Thus, we speculate that our results in the leukemic context could highlight a principle for elevated SG dependency that extends to other “stress prone” cancer types while also presenting a potential strategy to stratify patients for elevated therapeutic efficacy of SG targeting.

Towards elucidating the molecular functions of SGs in mediating AML/LSC stress adaptation we applied an approach that represents the first use of low input BioID that we are aware of to dissect the SG proteome in primary patient samples. Integrating these protein interaction networks with those at the RNA level also nucleated by G3BP1 and G3BP1-dependent transcriptional outcomes we reveal the G3BP1^+^ SG-mediated orchestration of a consolidated regulation of key AML- and LSC-supportive programs including LSC signatures, MYC targets, innate immune, inflammatory and apoptotic signaling. In particular, G3BP1 recognizes highly structured 3’UTRs within transcripts encoding pro-apoptotic effectors and, in coordination with its SG interactor UPF1, inhibits this subset of targets via structure-mediated decay. While sequestration of apoptotic proteins to SGs has been shown in other cell types and is apparent in our AML SG proteome as well, the mode of G3BP1 regional binding and targeted degradation of pro-apoptotic transcripts is a novel mechanism of SG-mediated survival. Fisher et al first presented coordination between G3BP1 and UPF1 to facilitate structure-mediated decay of target mRNAs but by sequencing transcripts associated with the insoluble SG solid-state core did not localize this mechanism to SGs^43^. Our contrasting use of eCLIP-seq provides us a more encompassing view of both the transcripts associated with the SG cores and their liquid-like shells, where more dynamic shuttling of proteins and RNA is known to occur^24,50^. In addition, the increased representation of transcripts with highly structured 3’UTRs in eCLIP datasets we observe following acute stress-induced enhancement of SGs supports the conclusion that this regulatory mechanism does indeed occur in AML SGs, and most likely within the liquid-like layer. Thus, while SGs have been traditionally viewed as hubs for stabilizing transcripts under stress, our work supports a novel emerging paradigm that through recruitment of factors including UPF and recently identified XRN proteins, regulated RNA decay is also localized to SGs^51^. These findings are provocative in suggesting the possibility that evolutionary pressures have selected for added structure in the 3’UTRs of transcripts that require careful control through stress. Moreover, it is intriguing to speculate that such an SG-mediated decay mechanism may also present a more pervasive phenomena in other cancer contexts.

Innate immune and inflammatory signatures undergo a similar consolidated regulation via SG-facilitated structure-mediated decay, as well as protein compartmentalization as we observed for apoptotic regulators. This finding is somewhat paradoxical to heightened inflammatory signaling being a hallmark of leukemia and transformation of HSCs^18^. However, key evidence now supports a paradigm shift that even this elevated inflammatory status must exist in a “Goldilocks” zone. For instance, in a mouse model of AML, animals treated with low levels of IFNγ exhibited reduced survival as AML growth was fueled by the mild inflammatory cues, high treatment doses on the other hand eliminated the AML disease and significantly improved survival^52^. More recently, Ellegast et al. have shown that cell-intrinsic inflammatory signals in AML blasts are under repressive control by IRF2BP2, and release of this repression induces excessive TNFα/NF-Kb signaling serving to trigger AML cell death^53^. Interestingly SGs have also been shown to control the balance of cell-intrinsic immune response to pathogen (dsRNA) stimulation^13^. Thus, it is intriguing to speculate that one of the contributions of SGs in AML is to maintain optimal levels of inflammatory signaling and that its inhibition unleashes a pro-death inflammatory response within leukemic cells. Such a mechanism may also be relevant in other cancers, where pro-apoptotic, inflammatory or heretofore unknown subclasses of cancer-type specific “stress transcripts” may operate under this non-traditional negative regulation via SGs to maintain their expression within the optimal tumor-supportive threshold.

While we have described an elevation of the SG machinery at the transcript level in AML LSCs the upstream regulators remain to be determined. One potential contributor is additional differential stressors experienced in this sub-compartment. Here it is intriguing to speculate that splicing dysregulation could be a factor. This comes in light of the recent demonstration that introduction of a mutation in the splice factor U2AF1, an event that defines a subset of MDS and AML patients, leads to elevated SG nucleation in response to oxidative stress and was linked to increasing the SG expression score potentially through altered binding and splicing of SG associated transcripts^54^. A similar connection was also found in an RNAi screen where knockdown of several proteins involved in RNA splicing, including U2AF1, resulted in the spontaneous formation of SGs in HeLa cells^55^. Together these studies implicate the mutation or altered expression of splice factors as regulators of SG dynamics. We find these observations particularly intriguing when considered with our recent study uncovering the heightened expression of many RNA splicing factors in the AML LSC compartment and specifically identifying the absolute requirement on RBM17 for LSC function^56^. This not only showcases that altered splicing can be particularly focused to the LSC compartment but considered alongside the abovementioned studies, seeds the idea that the altered splicing in LSCs in particular may synergize with background inflammation stress experienced in AML to prove a potent driver of SG elevation and dependency in this critical cell type.

While SGs have been implicated in solid tumors, a definitive exploration for any role of SGs in the cancer initiating cells that drive them has not been undertaken. By coupling state of the art in vivo xenotransplantation assays and imaging methodologies of patient samples, our work significantly forwards the paradigm that a strong, and in some cases preferential, reliance on elevated SG machinery and assembly sustains the cancer stem cell population in AML, underscoring a potentially overlooked contribution of SGs to other cancer stem cells pathobiology to initiation or seeding of other cancer types. Indeed, while SGs were not directly evaluated, hints that SGs may contribute to breast tumor initiating cells have been provided in a study that knocked down G3BP2, an SG nucleator and G3BP family member, and showed impaired mammosphere forming capacity^15^. Our findings thus present strong impetus to pursue the potential that SGs may be a crucial axis controlling stemness in cancer more generally. Future work interrogating this concept will require robust measures of stemness and renewal as it is clear from our analysis that inferring SG protein essentiality across cancer cell lines does not effectively highlight their potential importance in the rare but disease-causing cells.

Altogether our work advances a fundamental understanding of stress adaptive programs harnessed by LSCs for their persistence. We have described in high-resolution the AML-specific SG network, uncovering a thus far underappreciated mechanism of precision gene expression regulation via SG-mediated RNA decay. By these means we demonstrate that SGs are critical for restraining cell death responses in AML, control other critical and context-specific signatures in AML/LSCs, including inflammatory signaling, and represent novel targets for anti-AML therapeutic development.

## ACKNOWLEDGEMENTS

This work was supported by a Canadian Institutes of Health Research (CIHR) Doctoral Scholarship (A. Tajik and H.T. Chen), a Natural Sciences and Engineering Research Council of Canada (NSERC) Doctoral Scholarship (E. Tsao), an Ontario Graduate Scholarship (P. Joshi), a CIHR grant (#185952 to K.J. Hope), an NIH grant (#CA273432 to K.J. Hope. and G.W. Yeo) and an Ontario Institute for Cancer Research (OICR) Investigator Award (K.J. Hope).

## METHODS

### Mouse Maintenance and Transplants

B6.SJL (Ly5.1+, RRID: IMSR_JAX:002014), and NSG (RRID: IMSR_JAX:005557) and NSG-SGM3 (RRID: IMSR_JAX:013062) mice were bred and maintained at the University Health Network. All animal experiments were performed in accordance with institutional guidelines approved by the institutional Animal Research Ethics Boards. Twenty-four hours prior to transplantation by tail vein or intrafemoral injection, mice were sublethally irradiated (580 rad or 315 Rad). At endpoint, BM and spleen were harvested, crushed in RPMI + 10% FBS or IMDM + 2% FBS, and passed through 40-μm cell strainers (Corning; cat. no. 352340). Ammonium Chloride (STEMCELL Technologies; cat. no. 07850) was used for lysis of red blood cells.

### Lentivirus Production

Lentivirus was prepared by transient transfection of Lenti-X 293T (Cadarlane cat. no. 632180) cells with pMD2.G (RRID: Addgene_12259) and psPAX2 (RRID: Addgene_12260) packaging plasmids to create VSV-G pseudotyped lentiviral particles, as previously described^57^. All viral preparations were ultracentrifugated, resuspended in low volumes, and titered on HeLa cells (RRID: CVCL_0030) before being used to infect primary cells and cell lines.

### RN2c Murine Leukemia Transplantation

RN2c cells (MLL-AF9/NrasG12D/hCas9; a kind gift from Dr. Vakoc, Cold Spring Harbor Laboratory; received in 2015) were thawed and plated in RPMI supplemented with 10% FBS, at a maximum density of 3 million cells per mL. RN2c cells were all used at approximate passage number in vivo of 2–3. Cells were incubated for 1-2 hours to recover. 5 μg/mL polybrene (MilliporeSigma; cat. no. H9268) was added. Pretitrated lentivirus was added at a clonal MOI of 0.2 and cultures were incubated overnight, after which they were spun for 5 minutes at 1,500 rpm, resuspended in fresh RPMI supplemented with 0.5% FBS. The GFP+ fraction was determined 24 hours after transduction, and 1 million cells per mouse were intravenously transplanted (tail vein) into sublethally irradiated B6.SJL (Ly5.1, RRID: IMSR_JAX:002014) recipient mice. After 10 days (T10 primary), leukemic BM and splenocytes were isolated for subsequent flow analysis. Mouse BM was blocked with mouse BD Fc Block (BD Biosciences; cat. no. 553142, RRID: AB_394657). Cells were subsequently stained with fluorochrome-conjugated antibodies: CD45.2 V450 (BD Biosciences; cat. no. 560697, RRID: AB_1727495); and CD117 (c-Kit) APC (BD Biosciences; cat. no. 553356, AB_3985366) for quantitative analysis by flow cytometry.

### Flow Cytometry

All flow cytometry analysis was performed using a BD LSRII Analyzer, BD LSRFortessa, or BD LSRFortessa X-20 (BD Biosciences). Analysis was performed using FlowJo software (Tree Star, RRID: SCR_008520).

### Enrichment Analysis

The GSEA application (v4.3.2)^58,59^ was used to perform preranked GSEA against a gene set repository maintained by Dr. Gary Bader (Princess Margaret Cancer Centre), which encompasses gene sets from GO Biological Processes, Reactome, and MSigDB. Rank scores were calculated as −log10(P)*sign(log2FC). Stress granule proteomes were collected from Jain et al 2016^24^ and the RNAgranuleDB v2.0^21^.

LSC data was collected from RNA-sequencing data on 72 LSC+ (engrafting) and 38-LSC-(non-engrafting) AML fractions evaluated for functional LSC activity via xenotransplantation^60^.

Relapse data was collected from RNA-sequencing on 68 paired AML samples collected at diagnosis and relapse following induction chemotherapy, compiled from 1 in-house cohort (John Dick Lab, unpublished) and 4 publicly available cohorts^25,61–63^. Genomics data was collected from RNA-sequencing data for 1034 patients from 3 independent cohorts (TCGA, BEAT-AML, Leucegene). Gene expression was linked to genomics and clinical characteristics based on clinical data spanning 39 point mutations, 92 mutation combinations, 12 cytogenetic alterations and 9 FAB morphological classes.

### Intracellular Staining

OCI-AML22 cells were thawed and cultured as described previously (Boutzen et al. 2022)^27^. Approximately 100,000 cells were blocked with human IgG (MilliporeSigma; cat. no. I4506, RRID: AB_1163606). Cells were subsequently stained with fluorochrome-conjugated antibodies: CD34 APC (BD Biosciences; cat. no. 555824, RRID: AB_398614) and CD38 PE (BD Biosciences; cat. no. 347687) then fixed and permeabilized according to BD Cytofix/Cytoperm™ Fixation/Permeabilization Solution Kit (BD Biosciences: cat. no. 554714) guidelines. G3BP1 Rabbit polyclonal antibody (ProteinTech; cat. no. 13057-2-AP) was incubated overnight. Secondary staining was completed with Alexa Fluor® 488 Donkey Anti-Rabbit IgG (H+L) (ThermoFisher: cat. No A-21206).

### Immunofluorescence

Primary AML or cell lines were treated via heat shock (42-45 °C) for 20-60 minutes or vinorelbine (Millipore Sigma: cat. No V2264-5MG) (final concentration 200 µg/mL) for 1 hour. AML were prepared for staining by cytocentrifugation (200 xg, 10 minutes) into a well of u-Slide 18-well Glass Bottom plate (IBIDI: cat. No. 81817). Cells were fixed in 4% PFA (Electron Microscopy Sciences; cat. no. 15710) for 20 minutes, followed by permeabilization in 0.1× Triton X-100 (Bioshop; cat. no. TRX777) in PBS for 10 minutes. Samples were incubated in blocking buffer (PBST + 10% goat serum + 1% W/V BSA) for 1 hours at room temperature, followed by incubation with anti-G3BP1 (Proteintech; cat. no. 13057-2-AP) overnight at 4 °C. Secondary antibody incubation was performed with AlexaFluor-647 donkey-anti-mouse IgG (Thermo Fisher Scientific; cat. no. A21235) or AlexaFluor-488 donkey-anti-rabbit IgG (Thermo Fisher Scientific; cat. no. A21206) for 1 hour at room temperature with Hoechst 33342 Solution (20 mM) (Thermo Fisher Scientific; cat. no. 62249) at a 5 µM final concentration. Slides were mounted with Fluoromount mounting medium (Thermo Fisher Scientific; cat. no. 00-4958-02) and images were captured using a Widefield - Zeiss AxioObserver1 or Confocal - Leica SP8 (63× objective lens). Analysis was completed using the Cellprofiler application^64^. For primary AML analysis, stress granule analysis was completed in the cytoplasmic space over the nuclear region to avoid artifact identification in the periphery of cytospun cells. For Amnis ImageStream analysis, AML and CB were stained as described in the Intracellular staining protocol and imaged at 60x objective. Ideas Software was used for custom analysis and machine learning analysis of stress granule formation. For machine learning pipeline development of Live/Dead and SG+/SG- identification, a minimum of 30 manually selected photos for each condition were selected for pipeline development where Live/Dead determination was determined based on nuclear staining and brightfield images and SG+/SG- identification was based on G3BP1 staining.

### Live-Cell Imaging

For generation of the EGFP-G3BP1 fusion lentiviral vector the truncated-NGFR MA1 vector previously described in Rentas et al. (2016)^65^ was modified to first delete the mCMV-EGFP reporter region at EcoRV and NheI sites. The EGFP-linker-human G3BP1 was then cloned after the hPGK promoter replacing truncated-NGFR. Pre-titrated virus was used to infect THP-1 AML at an MOI of 2 to achieve >90% infection which remained stable in culture and with viably freezing cells. GFP-G3BP1 fusion THP-1 cells were incubated at 37 °C or heat shocked (42-45 °C) for 20- 60 minutes and plated in one well of u-Slide 18-well Glass Bottom plate (IBIDI: cat. No. 81817) for live cell imaging on the Widefield - Zeiss AxioObserver1 (63X objective).

### Culture of Primary AML Patient Samples

All AML patient samples were obtained as peripheral blood draws with written informed consent and conducted in accordance with recognized ethical guidelines by the Research Ethics Boards at University Health Network (UHN) Research Ethics Board (CAPCR # 20-6026) in accordance with Canadian Tri-Council Policy Statement on the Ethical Conduct for Research Involving Humans (TCPS). Immediately following harvest samples were subjected to Ficoll-Paque PREMIUM (Cytiva; cat. no. 17544203) separation, mononuclear cells were stored in the vapor phase of liquid nitrogen in 10% DMSO, 40% FCS and alpha MEM. Primary samples were thawed in X-VIVO (Lonza; cat. no. BEBP04-743Q) 50% FBS with 100 μg/mL DNAse prior to using in in vitro and in vivo assays. Primary AML samples were grown in AML growth media consisting of X-VIVO with 20% BIT Serum Substitute (STEMCELL Technologies; cat. no. 09500) or StemSpan SFEM II (STEMCELL Technologies; cat. no. 09655), supplemented with 100 ng/mL human SCF (R&D Systems; cat. no. 255-SC-050), 10 ng/mL human IL3 (R&D Systems; cat. no. 203-IL-050), 20 ng/mL human IL6 (PeproTech; cat. no. AF-200-06), 20 ng/mL human TPO (PeproTech; cat. no. AF-300-18), and 100 ng/mL human FLT3 L (R&D Systems; cat. no. 308-FKN-100).

### Lentivirus-Infected Primary AML Transplantation Assays

Production of shG3BP1 and shScramble expressing lentiviral particles was performed as previously described^57^ and briefly described under “Lentivirus Production” and validated by qRT-PCR and/or western blot. For knockdown experiments, AML cells were thawed and transduced at an MOI of 50 for 24 hours. All cells were transplanted intrafemorally into sublethally irradiated (315 Rad) NSG mice (RRID: IMSR_JAX:005557). Mice were sacrificed approximately 8 weeks after transplant, and BM from the right femur (site of injection) and remaining tibias, pelvis, and left femur were harvested along with spleens, crushed, filtered, and red blood cell lysed using ammonium chloride (STEMCELL Technologies; cat. no. 07850). Cell suspensions were blocked with mouse BD Fc Block (BD Biosciences; cat. no. 553142, RRID: AB_394657) and human IgG (MilliporeSigma; cat. no. I4506, RRID: AB_1163606), respectively. Cells were subsequently stained with fluorochrome-conjugated antibodies: CD45 BV421 (BD Biosciences; cat. no. 563879, RRID: AB_2744402); CD33 PE (BD Biosciences; cat. no. 347787, RRID: AB_400350); CD14 PE-Cy7 (BD Biosciences; cat. no. 561385, RRID: AB_10611732) or APC-H7 (BD Biosciences; cat. no. 561384, RRID: AB_10611720); CD11b BV605 (BD Biosciences; cat. no. 562721, RRID: AB_2737745); CD34 APC (BD Biosciences; cat. no. 555824, RRID: AB_398614) and 7AAD PerCP-Cy5.5 (BD Biosciences; cat. no. 559925, RRID: AB_2869266) for quantitative analysis by flow cytometry. For Secondary limiting dilution transplantation, BM from primary engrafted AML mice were thawed and stained for CD45 and CD33 as described above and sorted on the BD FACS ARIA Fusion for CD45+CD33+GFP+. Equal numbers of shScramble and shG3BP1-4 cells were transplanted into secondary NSG-SGM3 mouse recipients. At endpoint, mouse BM was isolated and analyzed as described above. ELDA software was used for analysis of stem cell frequency^66^. A 0.2% splenic engraftment cut-off was used for ELDA. High dose = 250,000 GFP sorted cells. Low dose = 50,000 GFP sorted cells. Log-fraction plot of the secondary limiting dilution transplant engraftment data shows the log-active cell fraction represented by the slope. The 95% confidence interval is represented with the dotted lines. A down-pointing triangle represents the data value with zero negative response at the shScramble high 250,000 dose whereas circles represent the remaining doses whereby there is a minimal of one negative response (no engraftment).

### Knockdown and Overexpression Lentivector Design

For knockdown experiments, sample and control (shScramble) target sequences were cloned downstream of U6 via AgeI and EcoRI in PLKO.1-TRC (Addgene Plasmid 10878227) in which Puro was replaced with GFP (PLKO.1-TRC-GFP). AML cells were thawed, transduced at an MOI of 50 for primary AML or MOI 0.5-4 for cell lines with lentivirus expressing pLKO.1-GFP- shScramble, shG3BP1-1(5′- GAAGGCGACCGACGAGATAAT-3′), shG3BP1-3 (5′- GATGCTCATGCCACGCTAAAT-3′), or shG3BP1-4 (5′- AGTGCGAGAACAACGAATAAA-3′) and cultured for up to 10 days. For G3BP1 knockdown rescue assays, DAP (5’- ACCTAAACCCACTGTGTTCAT-3’), APAF1(5’- GCCATGTCTATAAGTGTTGAA-3’), and BCL2L11 (5’-AGCCGAAGACCACCCACGAAT-3’) target sequences were cloned into a pLKO.1 lentivector with a BFP reporter. For G3BP1 overexpression assays Human G3BP1 and truncated-NGFR control were expressed in MA1 downstream of hPGK promoter and bidirectionally to minimal CMV driving GFP expression as previously described in Rentas et al. (2016)^65^.

For phenocopy overexpression assays, Luciferase control and human DAP, APAF1, and BCL2L11 were expressed in the pSMALB vector downstream of an SFFV promoter and bidirectionally to minimal CMV driving BFP expression. The cDNAs for human DAP (Horizon Discovery MGC cDNA, MHS6278-202829566) and BCL2L11 (Horizon Discovery MGC cDNA, MHS6278-202801513) were amplified using primers containing PacI and SalI restriction sites. The amplified products were digested and subsequently cloned into the pSMALB vector. For APAF1, the protein-coding region (CCDS9069.1) was synthesized by Integrated DNA Technologies (IDT) in three gBlock segments. The first segment was engineered with a PacI site at the start, and the last segment included a SalI site at the end. Each segment contains a 30-bp overlap with its adjacent fragment, facilitating full-length assembly via fusion PCR. The complete APAF1 sequence was initially cloned into pCR™Blunt II-TOPO™ vector through TOPO™ cloning (Invitrogen; cat. no. K280020), excised by restriction enzymes, and then subcloned into the pSMALB vector.

### AML Immunophenotyping and Apoptosis

At their corresponding time points, cells were stained for quantitative flow-cytometric analysis. For evaluation of knockdown experiments: CD14 APC-H7 (BD Biosciences; cat. no. 561384, RRID: AB_10611720); CD11b BV605 (BD Biosciences; cat. no. 562721, RRID: AB_2737745); and 7AAD (BD Biosciences; cat. no. 559925, RRID: AB_2869266). For all in vitro primary AML treated with G3Ib experiments, AML cells were thawed and cultured with 50/25/10 μmol/L G3Ib or equivalent DMSO (Fisher Scientific; cat. no. BP231-100) volume for 3 or 7 days. At their corresponding time points, cells were subsequently stained for quantitative flow-cytometric analysis: Annexin V AlexaFluor-647 (Innovative Research; cat. no. A23204, RRID: AB_2341149); CD14 APC-H7 (BD Biosciences; cat. no. 561384, RRID: AB_10611720); and CD11b BV605 (BD Biosciences; cat. no. 562721, RRID: AB_2737745). Cell counts were calculated by combining cells with ViaStain AOPI Staining Solution (ESBE; cat. No. NEX- CS2010625ML) for live-cell counts on the Nexcelom K2 Cellometer.

### Isolation of Human Cord Blood HSPCs

All human umbilical CB samples were obtained with written informed consent and conducted in accordance with recognized ethical guidelines by the Research Ethics Board at UHN (REB # 20- 6026) in accordance with Canadian Tri-Council Policy Statement on the Ethical Conduct for Research Involving Humans (TCPS). Freshly harvested CB samples were stored for a maximum of 3 days after collection at 4°C and then mononuclear cells were collected by centrifugation with Ficoll-Paque PREMIUM (Cytiva; cat. no. 17544203), followed by red blood cell lysis with Ammonium Chloride (STEMCELL Technologies; cat. no. 07850). Cells were subsequently stained with a cocktail of lineage-specific antibodies (CD2, CD3, CD11b, CD11c, CD14, CD16, CD19, CD24, CD56, CD61, CD66b, and GlyA; STEMCELL Technologies; cat. no. 19356) for negative selection of lineage-depleted (Lin−) cells using an EasySep immunomagnetic column (STEMCELL Technologies; cat. no. 18000). Cells were stored as Lin− cells in the vapor phase of liquid nitrogen in 10% DMSO + 90% FBS.

### RNA Isolation, cDNA Synthesis and Quantitative Real Time PCR (qRT-PCR)

For all qRT-PCR determinations total cellular RNA was isolated with TRIzol LS reagent (Thermo Fisher Scientific; cat. no. 10296028) or Monarch total RNA miniprep kit (NEB; cat no. T2010S) according to the manufacturer’s instructions and cDNA was synthesized using the qScript cDNA Synthesis Kit (Quanta Biosciences; cat. no. 95048-100). The mRNA content of samples compared by qRT-PCR was normalized based on the amplification of GAPDH or ACTB. qRT-PCR was done in triplicate with TaqMan® Fast Advanced Master Mix (Thermo Fisher Scientific; cat. no. 4444964) with gene-specific TaqMan probes (Thermo Fisher Scientific, FAM-MGB). qRT-PCR was also completed using the Luna Universal qPCR Master Mix (NEB; cat. no. M3003X) with gene specific primers.

### Actinomycin D treatment

THP-1 cells were treated with 10 μg/mL actinomycin D (Millipore Sigma; cat. no. A1410-2MG) over a 6-hour time course whereby cells were harvested at 0, 3, and 6 hours post-actinomycin D treatment. RT-qPCR was performed to quantify RNA changes.

### Western Blot

Immunoblotting was performed with anti-G3BP1 rabbit polyclonal antibody (Proteintech; cat. no. 13057-2-AP), anti-G3BP1 mouse monoclonal antibody (Proteintech; cat. no. 66486-1-Ig), anti-β-actin mouse monoclonal (MilliporeSigma; cat. no. A5441). Secondary antibodies used were IRDye 680RD goat anti-rabbit IgG (LI-COR Biosciences; cat. no. 926-68071, RRID: AB_10956166), IRDye 680RD goat anti-mouse IgG (LI-COR Biosciences; cat. no. 925-68070, RRID: AB_2651128), and IRDye 800CW goat anti-mouse IgG (LI-COR Biosciences; cat. no. 926-32210, RRID: AB_621842).

### Inhibitor Assays

G3Ib was provided as a kind gift from Dr. Paul Taylor (PharmaResources). THP-1 EGFP-G3BP1 fusion expressing cells were thawed and expanded in RPMI supplemented with 10% FBS. Cells were then pre-incubated with G3Ib or Resveratrol (Millipore Sigma; cat. no. R5010-100MG) for 20 minutes and 1 hour respectively according to previously published data^32,33^. Cells were then stress treated via heat shock (43 °C) for 40 minutes or vinorelbine (Millipore Sigma: cat. no V2264-5MG) (final concentration 200 μg/mL) for up to 2.5 hours and imaged on the Widefield - Zeiss AxioObserver1 (63X objective). G3Ib treated cells were analyzed with a custom cellprofiler pipeline for SG determination and RSVL treated cells were analyzed via visually (blinded to conditions). For colony forming unit analysis, AML and CB cells were pre-treated with 50 μM G3Ib (or DMSO control) for 3 days prior to washing and counting live cells for plating equal number into MethoCult™ Enriched gel (STEMCELL Technologies; cat. no. 04435).

### RNA-seq in Primary AML

Five human primary AML samples were stained for CD34-APC (BD Biosciences; cat. no. 555824, RRID: AB_398614) and sorted on the BD FACS ARIA Fusion (BD Biosciences; Princess Margaret Flow facility). Cells were then infected with pLKO.1-EGFP-shScramble or-shG3BP1-4 at an MOI of 50 for 5 days. After 5 days, 7AAD-EGFP+ primary AML cells were then sorted (minimum 20,000 cells and RNA was isolated via PicoPure RNA Isolation Kit (Thermo Fisher Scientific; cat. No. KIT0204) following the manufacturer’s protocol, with the addition of a DNAse treatment (Qiagen: cat. no. 79254). Libraries were prepared using the Illumina Stranded mRNA Prep kit (Illumina; cat. no. 20040534) according to manufacturer instructions and sequencing was performed on the NextSeq2000 using the paired-end 100 cycle configuration at a depth of 20 million reads per sample. Short-read quality control was performed using *FastQC v0.11.5*^67^ and *MultiQC v1.7*^68^. Low-quality reads and the sequencing adapters were trimmed using *Trim Galore v0.6.6*. Subsequently, reads were aligned to GENCODE human reference genome *v38*^69^ using *STAR v.2.7.9a*^70^ and low-quality alignments (*mapq score* < 15) were filtered using samtools v1.17. Next, *featureCounts v2.0.1*^71^ was used to quantify the gene expression from alignment files. *DESeq2 v1.40.1*^72^ R package was used to identify the differentially expressed genes between treatment groups. Gene set enrichment analysis (GSEA)^73^ was performed using *fgsea v1.26.0*^74^ and the curated list of pathways published by Bader lab (http://baderlab.org/GeneSets - updated August 2022)^75^. All p-values were adjusted using the Benjamini-Hochberg method^76^ where applicable.

### RNA-seq and Proteomics in THP-1 AML

THP-1 were infected with pLKO.1-EGFP-shScramble or-shG3BP1-1/3/4 in triplicate at an MOI of 3.5. 7AAD-EGFP+ cells were sorted on day 4, replated and RNA isolated on day 5 via Monarch total RNA miniprep kit (NEB; cat no. T2010S) according to manufacture protocols. The KAPA Stranded RNA-Seq Kit (code KK8401) was used for library preparation and sequencing was performed on the Novaseq 6000 at a depth of 20 million reads per sample. Sequencing reads were checked for quality, aligned, and analyzed as described in ‘RNA-seq in Primary AML’.

For proteomic analysis, 500,000 cells were isolated on day 6, washed with phosphate-buffered saline (PBS) in triplicate, and subsequently resuspended in SP3 lysis buffer (100 mM HEPES, pH 8.0, 1% SDS) prior to flash freezing. The samples were then heated at 95°C for 10 minutes to denature proteins. To shear genomic DNA and facilitate further protein lysis, samples were subjected to sonication using a probe-less ultrasonic sonicator (Hielscher VialTweeter) with five 10-second cycles at 10 watts per tube. A 50 μg aliquot of the resulting protein lysate was used for downstream sample preparation.

Reduction of disulfide bonds was achieved by incubating the lysate with 5 mM dithiothreitol (DTT) for 30 minutes at 60°C. Free sulfhydryl groups were alkylated by treatment with 25 mM iodoacetamide (Sigma-Aldrich; cat. no. I1149) in the dark for 30 minutes at room temperature. Proteins were then captured using the SP3 bead-based protocol^77^. Magnetic beads (GE Healthcare; cat. no. 45152105050250) were added to the protein mixture in a 10:1 (w/w) ratio, and absolute ethanol was added to achieve a final ethanol concentration of 70%. The mixture was shaken at room temperature for 5 minutes at 1000 rpm, and the supernatant was discarded. The beads were washed twice with 80% ethanol, and the washes were discarded.

Protein digestion was performed in 100 mM ammonium bicarbonate buffer containing 2 μg of trypsin/Lys-C enzyme mix (Promega; cat. no. V5073) at 37°C overnight. The resulting peptides were desalted using C18-based solid-phase extraction (SPE) and lyophilized in a SpeedVac vacuum concentrator. Peptides were reconstituted in mass spectrometry-grade water containing 0.1% formic acid.

Peptides (2 μg) were analyzed using a two-column setup consisting of a 2 cm Acclaim PepMap 10 μm trap column (75 μm, 3 μm, 100 Å) and a 50 cm EasySpray ES803 column (75 μm, 2 μm, 100 Å) (Thermo Fisher Scientific) connected to an Easy nLC 1000 (Thermo Fisher Scientific) nano-flow liquid chromatography system, coupled to a Q Exactive mass spectrometer (Thermo Fisher Scientific). Peptide separation was achieved by reverse-phase chromatography using a 265-minute non-linear gradient (4-48% buffer B, 0.1% formic acid in acetonitrile) at a flow rate of 250 nL/min, with the column temperature maintained at 45°C. Data acquisition was performed in positive-ion mode with data-dependent acquisition (DDA) using a top-25 method. Full MS1 spectra were acquired in the m/z range of 350–1800 at a resolution of 140,000, with an automatic gain control (AGC) target of 3 × 10^6^ and a maximum ion fill time of 220 ms. MS/MS spectra were acquired at a resolution of 17,500, with an AGC target of 5 × 10^5^ and a maximum fill time of 45 ms. The isolation window width was set to 2.0 m/z, the isolation offset to 0.4 m/z, and the intensity threshold to 1.8 × 10^3^.

The raw data files were analyzed using MaxQuant software, with the UniProt human protein sequence database and a false discovery rate (FDR) threshold of 1% for peptide spectral matches. Search parameters included a maximum of two missed cleavages, oxidation of methionine as a variable modification, and carbamidomethylation of cysteine as a fixed modification. Intensity-based absolute quantification (iBAQ) and label-free quantification (LFQ) were enabled, with the match-between-runs feature disabled. Subsequent analyses were conducted using the proteinGroups.txt file, with contaminant sequences and decoy hits removed. Only proteins identified by two or more unique peptides were included in the final analysis.

### Enhanced Cross-Linking and Immunoprecipitation

Approximately 20 × 10^6^ THP-1 (or primary AML) cells per sample were heat shocked at 43 °C for 40 minutes (or 20 minutes for primary AML) and subsequently washed in PBS and UV-crosslinked at 400 mJ/cm2 on ice. Samples were then pelleted, snap-frozen, and stored at −80 °C. Enhanced Cross-linking and Immunoprecipitation (eCLIP) sequencing was performed as previously described^78^. Briefly, pellets were lysed in iCLIP lysis buffer, treated with RNase I (NEB; cat. no. M0314L) and Turbo DNAse (Thermo Fisher Scientific; cat. no. AM2239) for 5 minutes at 37 °C, followed by immunoprecipitation using 10 μg anti-G3BP1 Rabbit polyclonal antibody (ProteinTech; cat. no.13057-2-AP) and 125 μL M-280 Sheep-α-Rabbit IgG Dynabeads (Thermo Fisher Scientific; cat. no. 11203D) per sample. After several rounds of washing, samples were dephosphorylated (FastAP, Thermo Fisher Scientific; cat. no. EF0654; T4 PNK, NEB; cat. no. M0201L) followed by 3′ ligation (on-bead) with a barcoded RNA adapter followed using T4 RNA Ligase (NEB; cat. no. M0437M). Samples were again stringently washed, run on 4-12% Bis Tris gels, and transferred to nitrocellulose membranes. The region spanning 53–130 kDa was then isolated, followed by extraction of RNA from the membranes, and reverse transcribed (SuperScript III Reverse Transcriptase, ThermoFisher Scientific; cat. no. 18080044). A DNA adapter containing a 5′ random-mer was then ligated (3′), followed by the cleanup of the samples and PCR amplification. Libraries were size-selected using ProNex® Size-Selective Purification System (Promega; cat. no. NG2001) followed by sequencing on the the NextSeq2000 using the paired-end 100 cycle configuration. Reads were processed using the eCLIP processing pipeline for paired-end data, summarized as followed using the Skipper processing pipeline, available at https://github.com/YeoLab/skipper^79^. In summary, Skewer was used to trim reads of adapters^80^. Reads were then mapped with STAR (2.7.10a_alpha_220314)^81^ and PCR duplicates were removed with UMIcollapse^82^. A tiled window method was used for determining bound regions where reads were summed across evenly sized windows over the various genic regions and binned with consideration for GC biases. IP reads were then compared to size-matched input peaks to calculate enrichment of signal above input control (qmax < 0.05).

### BioID cell Line Generation and Processing

For generation of the miniTurbo-G3BP1 fusion lentiviral vector the truncated-NGFR MA1 vector previously described in Rentas et al. (2016)^65^ was modified. The MiniTurbo-3xFLAG-G3BP1 sequence was a kind gift from Dr. Payman Tehrani from Dr. Anne-Claude’s lab^83^. MiniTurbo-3xFLAG-G3BP1 was cloned after the hPGK promoter replacing truncated-NGFR. A Control vector was created by inserting a 2xP2A self-cleavage sequence after miniTurbo to allow for separate expression of miniTurbo and G3BP1. Pre-titrated virus was used to infect THP-1 AML, MOLM-13 AML, OCI-AML22 cells and patient AML using up to an MOI of 50. For CD34+ AML, cells were sorted for CD34 APC prior to transduction. Cells were then incubated in biotin-free media (with the exception of MOLM-13 cells) for 5-7 days in biotin-free media. For Biotin-free media preparation, Streptavidin Sepharose High Performance beads (Cedarlane; cat. no. 17-5113-01) were washed 3x in PBS and then incubated with media at 4 °C for a minimum of 1 hour prior to Steriflip-HV (Thermo Fisher Scientific; cat. no. SE1M003M00), 0.45 µm, filtration removal of beads. After cell incubation, cells were treated with 50 μM D-Biotin (Cedarlane; cat. no. BB0078-5G) for 0.5-2.5 hours and were GFP+ sorted, PBS washed, and flash frozen prior to storage at-80 °C. OCI-AML22 cells were sorted for CD34/CD38 fractions in addition to GFP sorting. For THP-1 sample preparation, samples were highly infected (>95% GFP+) and not sorted prior to freezing. THP-1 pellets were lysed in 1x RIPA buffer supplemented with Benzonase (Millipore Sigma; cat. no. 70664-3) followed by sonication and incubation with Sepharose High Performance beads for 3 hours at 4 °C. Beads were then wash in 50 mM ammonium bicarbonate and incubated with sequencing grade Trypsin (Fisher Scientific; cat. no. PR-V5113) for 16 hours followed by a 1 μg top-up of trypsin for an additional 2 hours. Beads were pelleted and the supernatant carrying bound peptides were speed vacuumed until dry and stored at-80 °C. The THP-1 samples were run on a Thermo Scientific Q Exactive HF hybrid quadrupole-Orbitrap mass spectrometer (LC-MS/MS System).

The remaining low input samples were processed as follows. Samples were lysed using 0.5% SDS RIPA lysis buffer by adding 80 μL to each sorted sample in PCR tubes. The tubes were sonicated in a bath sonicator for 5 minutes. To each sample, 100 ng of RNAase (Bio Basic; cat. no. RB0474) and 25 units of Benzonase (MilliporeSigma; cat. no. 71205-3) were added and the samples were end-over-end rotated for 30 minutes at 4 °C. Afterwards, samples were centrifuged at 14,000g for 20 minutes at 4 °C and the supernatant was collected to a fresh PCR tube. Each sample then received 5 μL of pre-washed (in RIPA wash buffer) MagReSyn Streptavidin MS (ReSyn BioSciences MR-STP) beads diluted 3x (15 μL total). Samples were incubated with end-over-end rotation for 3 hours at 4 °C. Then, the supernatant was removed to waste, and the beads were washed with 35 μL of 2% SDS wash buffer, 40 μL of RIPA wash buffer, 45 μL of RIPA wash buffer and 50 μL of TNNE wash buffer. Finally, the beads were solvent exchanged to 100 mM ammonium bicarbonate over three separate washes. The supernatant was removed from the magnetic beads and to each sample 10 μL of 100 mM ammonium bicarbonate containing 5 ng of trypsin and 5 ng Lys-C (Promega; cat. no. V5071) was added directly to the bead bed. Samples were then placed on a thermomixer with heated lid for overnight digestion at 37 °C with 1500 rpm mixing. Afterwards, an additional 5 μL of digestion buffer containing 2.5 ng of trypsin and 2.5 ng Lys-C was added with digestion continuing for an additional 3 hours. Then, digestion was quenched by the addition of 2 μL of formic acid, vortexing, and quick centrifugation. The supernatant is collected into an autosampler vial insert. The beads are then washed two times with pure water and each wash is combined in the insert.

### BioID Mass Spectrometry Analysis

Samples are loaded onto an Optimize Technologies (Oregon City, OR, USA) EXP2 Stem Trap cartridge trap contain 2.7 um ES-C18 trap material with a total 0.17 μL volumne in an external 6- port valve by a Vanquish Neo nano-LC operating in a back flushing trap-and-elute configuration. The valve and LC were controlled directly through Chromeleon 7 software. Peptides were then separated on a 60-minute gradient of 80% acetonitrile and 0.1% formic acid up to 42% composition of the mobile phase on either a home-packed silica column (50cm, 75 um ID, 1.9 um C18 particles, 200 nL/min flow rate) or an IonOpticks Aurora Ultimate column (25 cm, 75 um ID, 1.7 um C18 particles, 400 nL/min flowrate). The LC was coupled to a Bruker timsTOF SCP (Bruker, Billerica, MS, USA) operating in DDA-PASEF and DIA-PASEF. For both methods, ion mobility ranged from 0.65 to 1.3 1/K0 in 166 ms, with matching accumulation time and collision energy was tied to the mobility of an ion across a linear ramp from 20 eV at 0.6 1/K0 to 54 eV at 1.3 1/K0. The DDA-PASEF method consisted of 6 PASEF ramps for an approximate 1.2 s cycle time and ions were selected for MS/MS fragmentation via a polygonal filter. The DIA-PASEF method consisted of 29 m/z windows across a total m/z range of 300 to 1100 with a 1.0 m/z overlap between windows. The approximate cycle time was 1.9 s. DIA data files were first converted to the Spectronaut.htrms format using the Spectronaut HTRMS converter before being searched by Spectronaut 18.2.230802 (Biognosys, Schlieren, Switzerland) with the directDIA+ algorithm. The complete human proteome in SwissProt format (from Uniprot) with common contaminating proteins^84^ and 10 biotinylation enzymes (BirA*, TurboID, miniTurbo, UltraID, BASU, microID, microID2 lbmicroID2, AirID, and BioID2) was used as the search database. Search and validation parameters were the Spectronaut defaults. Precursors identified in less than 50% of runs were filtered and missing values were imputed using Spectronaut’s Wise Imputing method and all samples were run with cross-run normalization. Differential abundance was determined by unpaired t-tests with multiple testing correction.

DDA files were uploaded to a local ProHits LIMS^85^ for archiving and searching via an integrated implementation of FragPipe and MSFragger (v19)^86–88^. The complete human proteome in SwissProt format (from Uniprot) with common contaminating proteins^89^ and 10 biotinylation enzymes (BirA*, TurboID, miniTurbo, UltraID, BASU, microID, microID2 lbmicroID2, AirID, and BioID2) was used as the search database. MSFragger parameters were derived from the ‘default’ workflow option in Fragpipe except the isotope error was 0, and cysteine acetylation was disabled. Files were searched in parallel with separate calibration and optimization performed on each file individually. Results were combined and validated using MSbooster, Percolator, and Philosopher via Fragpipe with default settings except razor peptides were included for FDR scoring. SAINTexpress analysis was performed on SPC values using CRAPome (https://reprint-apms.org). Enrichment maps and cluster labels were generated using Cytoscape software (v.3.8.2) and EnrichmentMap (v3.0 and 3.3.1) and AutoAnnotate (v1.3.4) apps. Dotplots were generated on https://prohits-viz.org/.

---

**Supplementary Figure 1.**
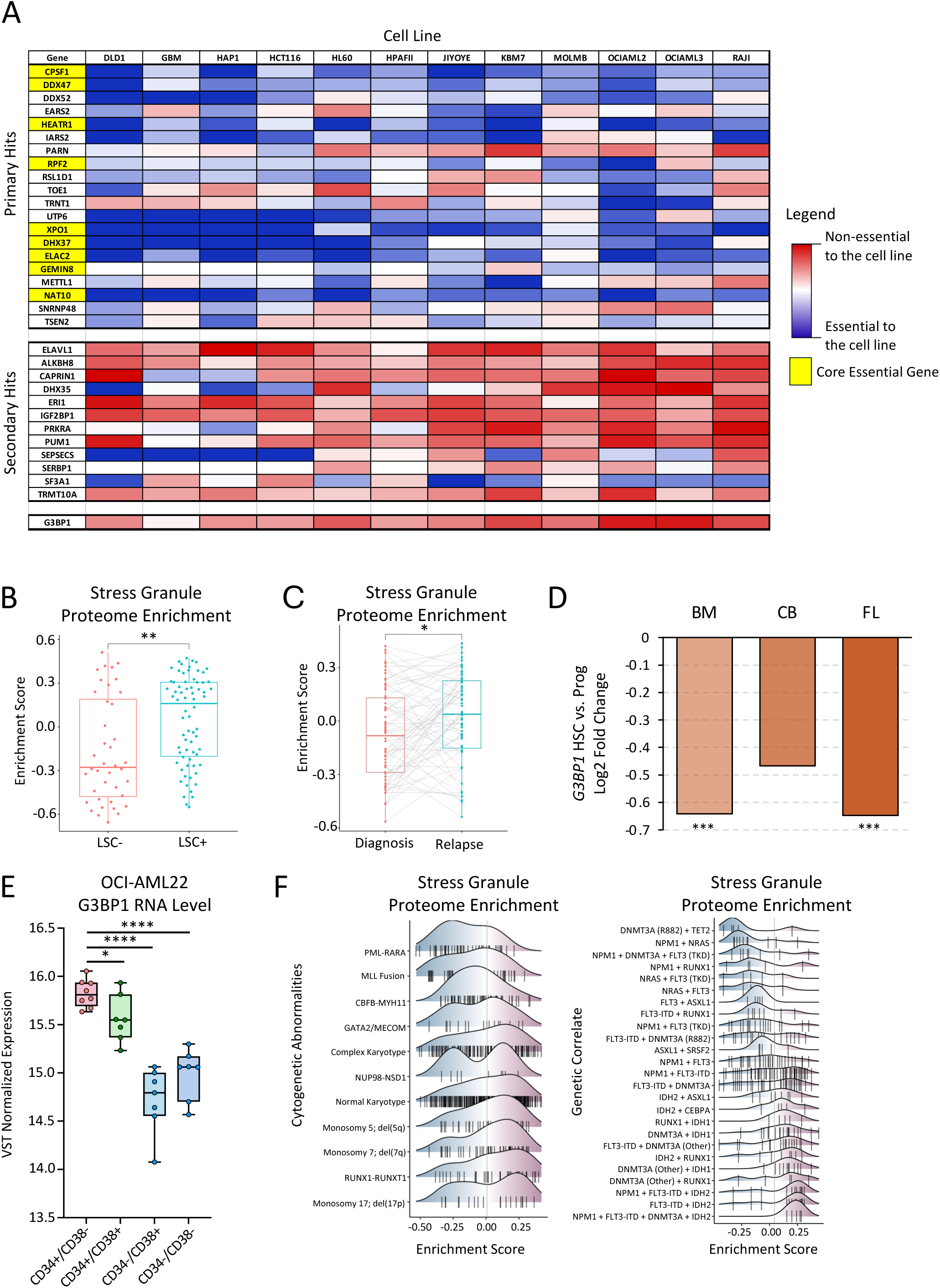
A) Heatmap of primary and secondary screen hits general essentiality score^22^ across various cell lines. RBPs highlighted in yellow are common essential genes and are enriched amongst primary hits versus secondary hits. **B-C)** The Tier 1 SG proteome^21^ is transcriptionally enriched in functionally validated LSC population^60^ **B)** and patient relapse vs diagnosis samples **C). D)** Relative RNA expression changes (log2 fold change) of G3BP1 in human fetal liver, cord blood and bone marrow across HSCs (CD34^+^CD38^-^CD90^+^CD45RA^-^) and progenitor cells (CD34^+^CD38^+^)^26^. **E)** RNA expression of G3BP1 in OCI-AML22 stem (CD34^+^CD38^-^), progenitor (CD34^+^CD38^+^) and non-stem (CD34^-^) cells. **F)** Density plots depicting Tier 1 SG proteome^21^ expression correlation analysis according to genomic abnormalities (right) and cytogenetic abnormalities (left). \*\*\*\**P* <.0001, \*\*\**P* <.001, \*\**P* <.01, \**P* <.05. n.s. not significant.

**Supplementary Figure 2.**
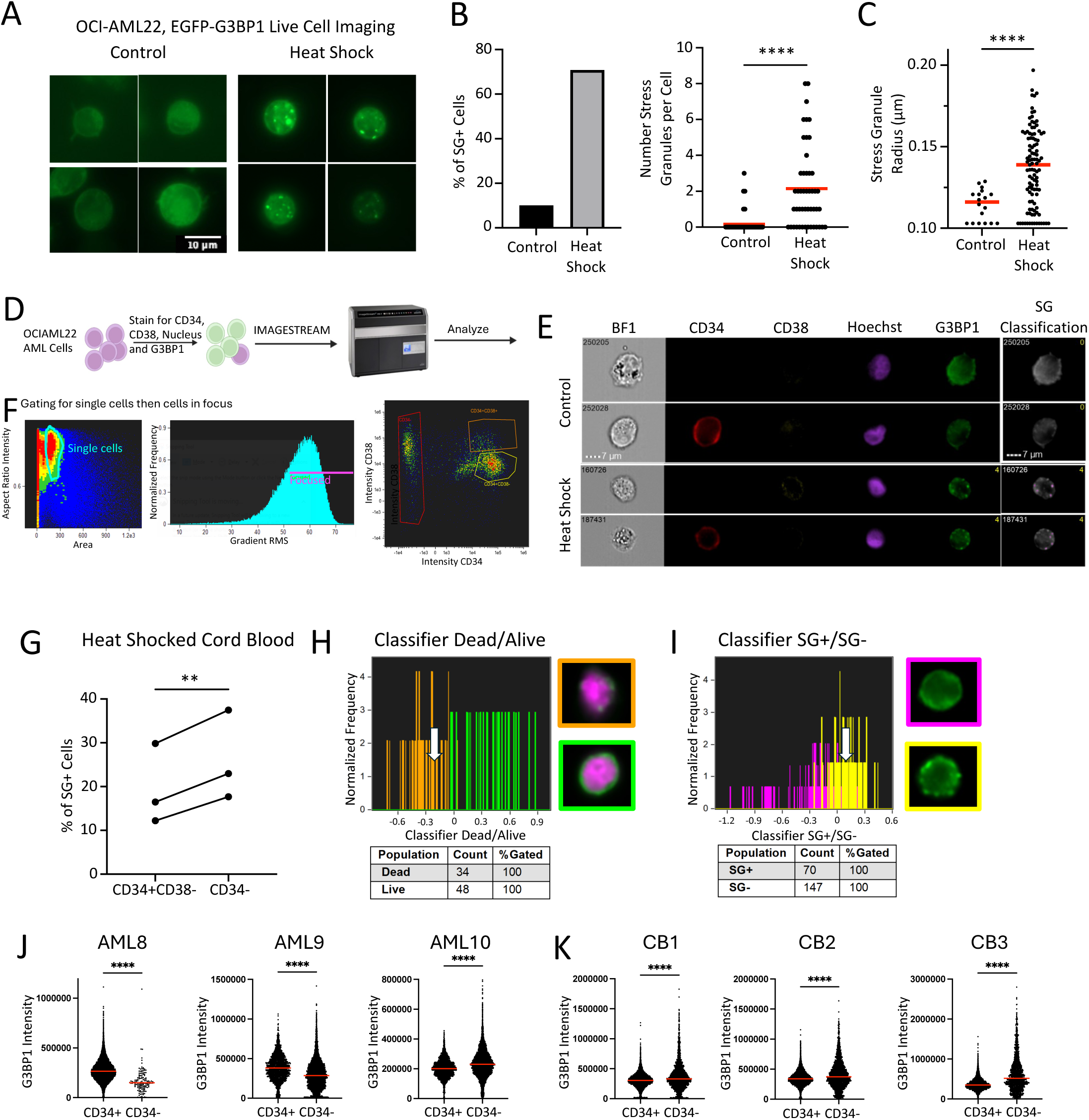
A) Representative images of live-cell EGFP-G3BP1 OCI-AML22 cells under heat shock and unstressed conditions. **B-C)** Cellprofiler based quantification of SGs **B)** and average SG radius **C)** in untreated and heat shocked OCI-AML22 EGFP-G3BP1 cells. **D)** Schematic of the Imagestream sample preparation pipeline. **E)** Representative Imagestream images of stressed (heat shock) and unstressed OCI-AML22 cells. **F)** Representative Imagestream gating of live/single OCI-AML22 cells followed by gating of cells in focus andCD34/CD38 fractions. **G)** Imagestream analysis of percent SG^+^ cells within the CD34^+^CD38^-^ and CD34^-^ fractions of 3 heat shocked biological CB replicates. **H)** Machine learning based separation of human live (green) vs dead cells/debris (orange). A cut-off of-0.2 was selected for primary AML and CB analysis. **I)** Machine learning based separation of human SG^+^ (yellow) vs SG^-^ (pink) cells. A cut-off of 0.1 was selected for primary AML and CB analysis. **J-K)** Imagestream analysis of relative total (punctate and diffuse) G3BP1 expression in 3 biological replicates of unstressed CD34^+^ or CD34^-^ AML cells **J)** or CB cells **K)**. \*\*\*\**P* <.0001, \*\*\**P* <.001, \*\**P* <.01, \**P* <.05. n.s. not significant.

**Supplementary Figure 3.**
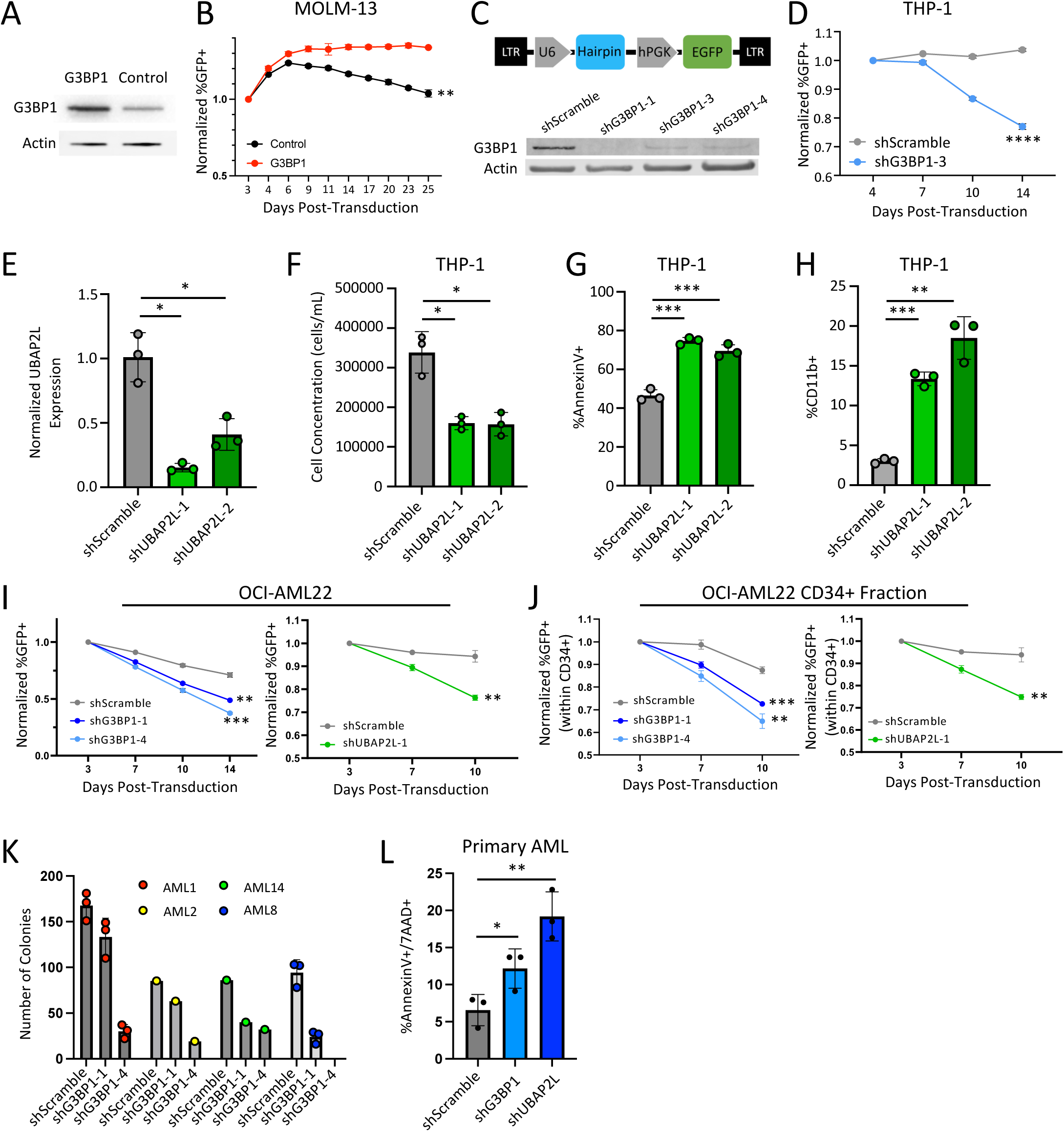
A) Western blot validation of G3BP1 overexpression. **B)** G3BP1 overexpression competitive cultures of MOLM-13 AML. **C)** Lentivector schematic for short hairpin knockdown in the PLKO.1 vector (top) and western blot validation of G3BP1 knockdown (bottom). **D)** G3BP1 knockdown THP-1 cell competitive growth cultures. **E)** qPCR validation of UBAP2L knockdown. **F-H)** UBAP2L knockdown THP-1 cell competitive growth in culture **F)**, increased apoptosis **G)**, and differentiation marked by CD11b **H)**. **I-J)** G3BP1 (left) and UBAP2L (right) knockdown OCI- AML22 total cell **I)** or CD34+ LSPC cell **J)** competitiveness in culture. **K)** Individual patient AML colony forming unit assays upon G3BP1 knockdown for four primary AML replicates in Figure 3G. **L)** Apoptosis in shG3BP1 and shUBAP2L primary AML16. \*\*\*\**P* <.0001, \*\*\**P* <.001, \*\**P* <.01, \**P* <.05. n.s. not significant.

**Supplementary Figure 4.**
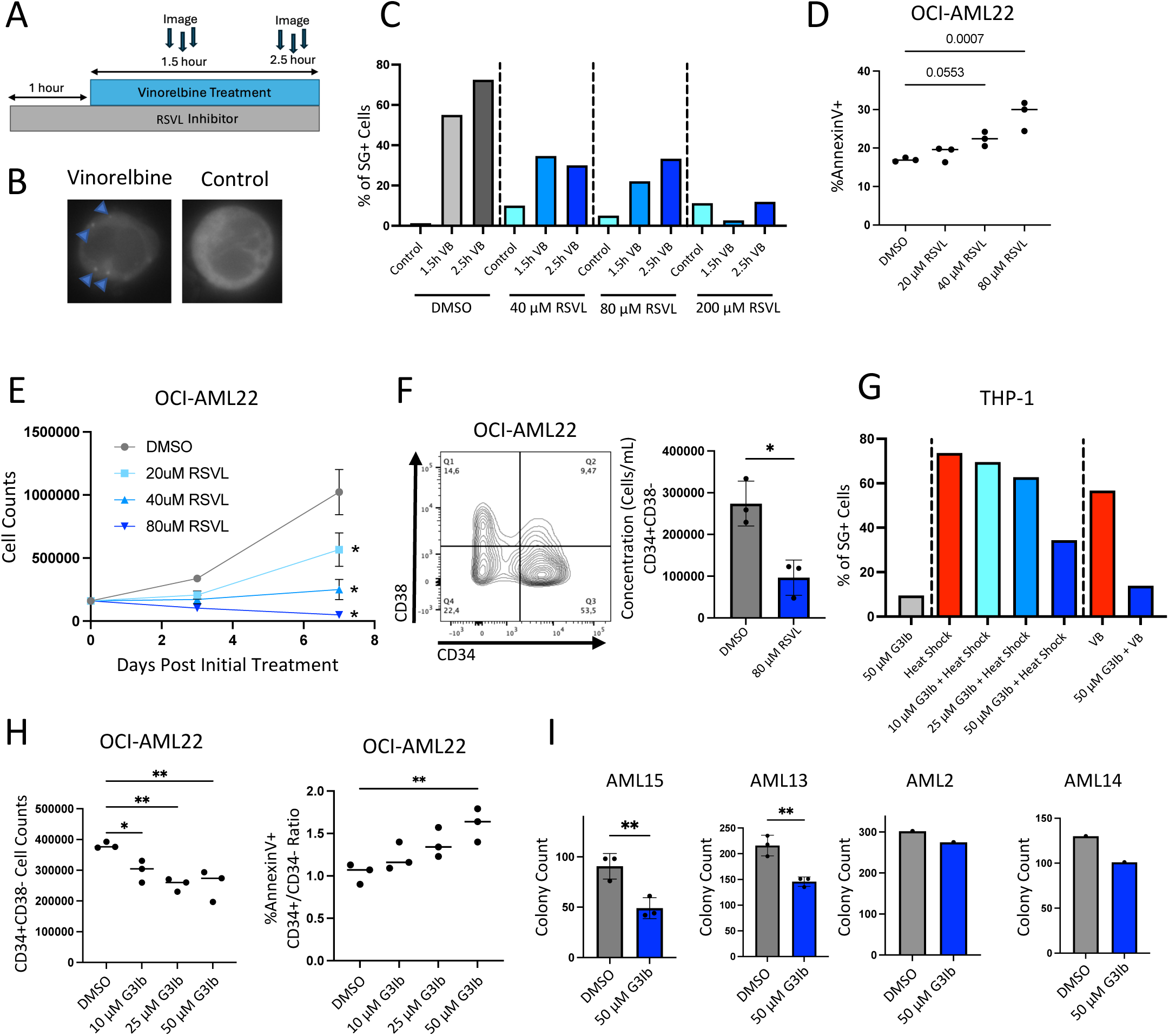
A) Schematic of RSVL pretreatment of THP-1 EGFP-G3BP1 cells followed by vinorelbine stress (100 μg/mL). **B)** Representative images of vinorelbine induced SGs in GFP-G3BP1 THP-1 cells. **C)** Percent of THP-1 EGFP-G3BP1 cells pre-treated with resveratrol prior to vinorelbine stress (100 μg/mL) for 1.5 or 2.5 hours that formed SGs. **D)** Apoptosis of OCI-AML22 cells treated for 3 days with either 20 μM, 40 μM, or 80 μM RSVL. **E)** OCI-AML22 cell fold changes in culture following RSVL treatment. **F)** Representative flow plot and changes in OCI-AML22 CD34+CD38- LSPC concentration following 7 days of RSVL treatment. **G)** Percent of THP-1 EGFP-G3BP1 cells pre-treated with G3Ib prior to heat shock or vinorelbine stress that formed SGs. **H)** CD34+CD38- OCI-AML22 cell counts (left) and apoptosis measurements in the CD34+ fraction (right) after 7 days of culture with 0-50 μM G3Ib treatment. **I)** Individual primary AML CFU assays upon G3Ib treatment. \*\*\*\**P* <.0001, \*\*\**P* <.001, \*\**P* <.01, \**P* <.05. n.s., not significant.

**Supplementary Figure 5.**
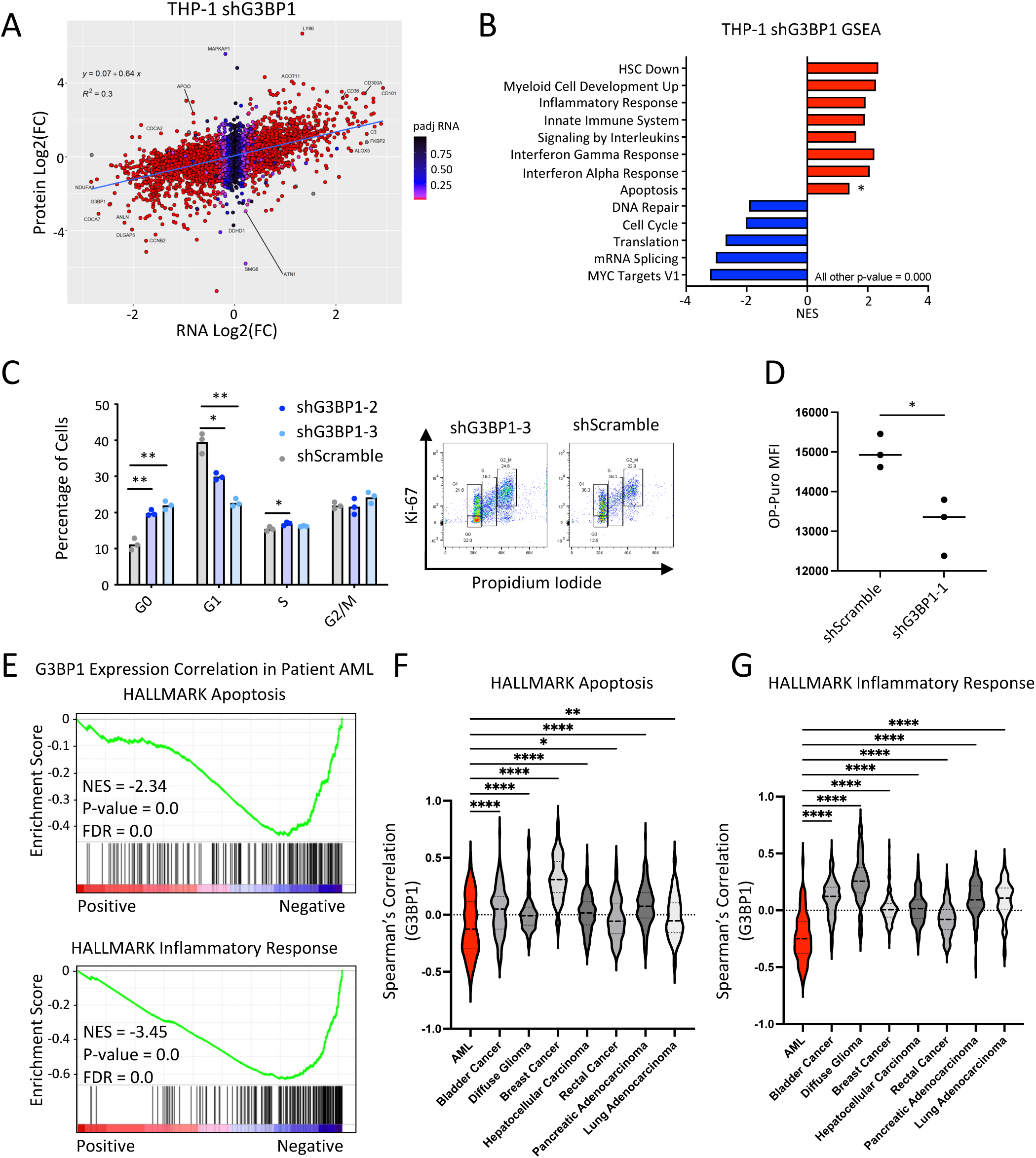
A) Scatterplot of transcript and protein expression, measured by RNA-seq and MS respectively, in shG3BP1-4 knockdown THP-1 cells. **B)** Barplot showing notable positive and negative enrichments with G3BP1 knockdown RNA-seq analysis in THP-1. **C)** Cell cycle assessment of G3BP1 knockdown THP-1 cells by Ki-67 and propidium iodine staining and flow cytometric evaluation including representative flow plots (right). **D)** Global protein synthesis in G3BP1 knockdown THP-1 cells measured by OP-Puro assay. **E)** GSEA of Patient AML G3BP1 gene expression correlation analysis shows G3BP1 expression is anticorrelated with genes in Hallmark apoptosis or inflammatory response. **F&G)** Average Spearman’s correlation coefficient of genes belonging to Hallmark Apoptosis **F)** or Hallmark Inflammatory Response **G)** gene sets with G3BP1 levels across various cancers. \*\*\*\**P* <.0001, \*\*\**P* <.001, \*\**P* <.01, \**P* <.05. n.s. not significant.

**Supplementary Figure 6.**
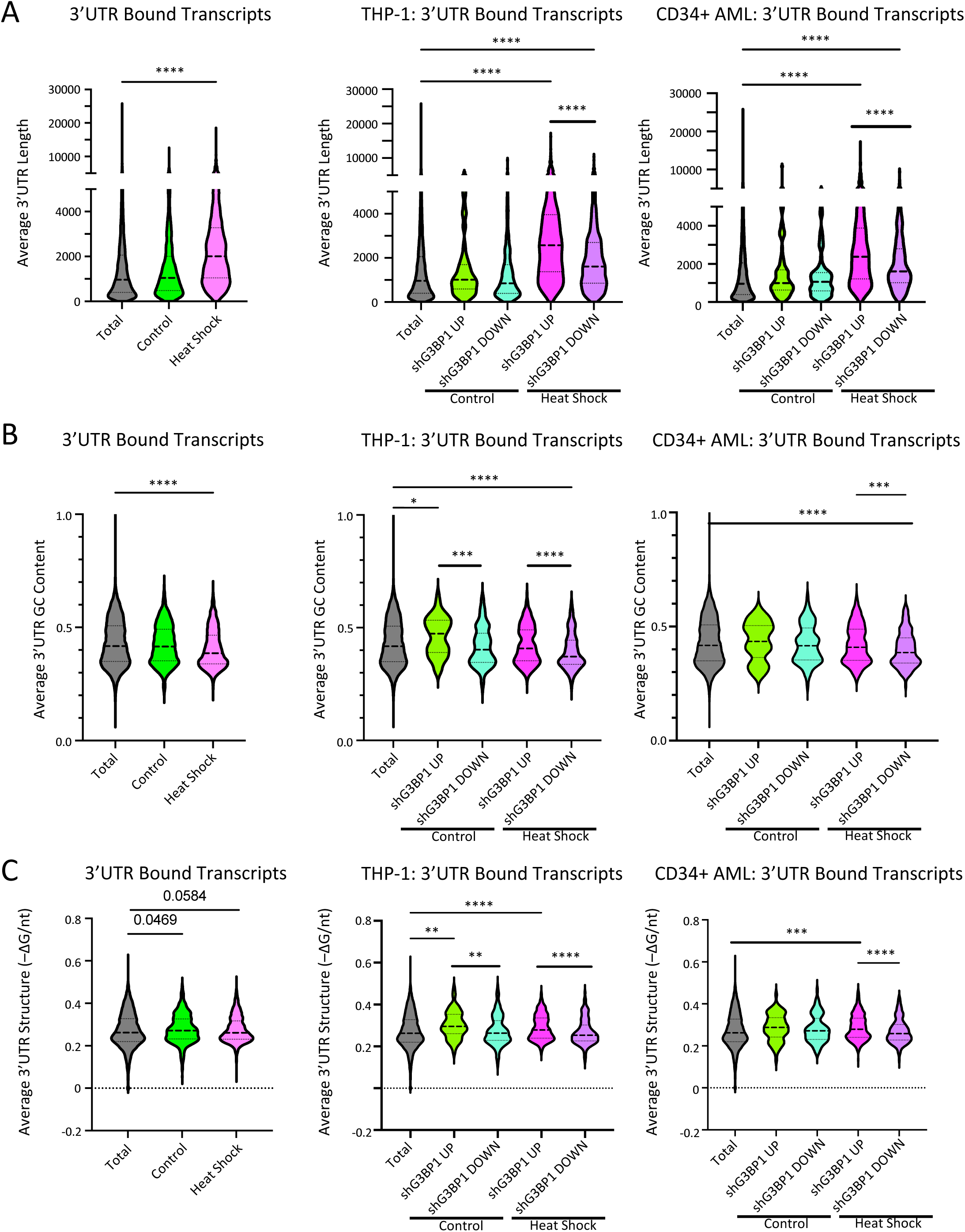
A) Average 3’UTR length of eCLIP-identified G3BP1 bound transcripts in untreated versus heat shocked THP-1 cells (left), further categorized by differential expression status upon G3BP1 knockdown in THP-1 (middle) and primary AML (right). **B)** Analysis of the average 3’UTR GC content of eCLIP identified G3BP1 bound transcripts in untreated versus heat shocked THP-1 cells (left), further categorized by differential expression status upon G3BP1 knockdown in THP-1 (middle) and primary AML (right). **C)** Analysis of the average 3’UTR structure of eCLIP identified G3BP1 bound transcripts in untreated versus heat shocked THP-1 cells (left), further categorized by differential expression status upon G3BP1 knockdown in THP-1 (middle) and primary AML (right). \*\*\*\**P* <.0001, \*\*\**P* <.001, \*\**P* <.01, \**P* <.05. n.s., not significant.

**Supplementary Figure 7.**
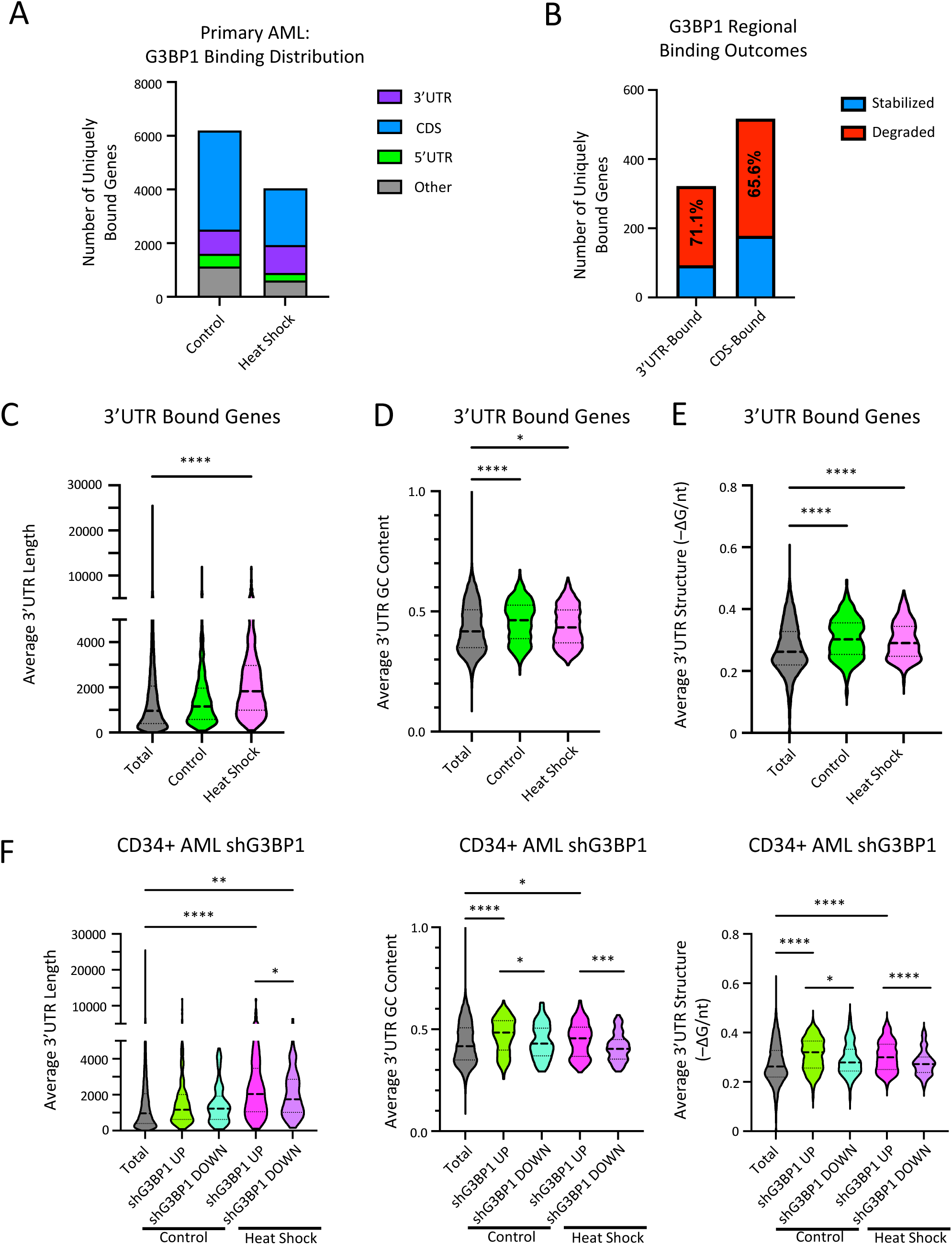
G3BP1 SG+ 3’ UTR transcriptome profiles in Primary AML **A)** Comparison of G3BP1-eCLIP binding distribution in untreated vs heat shocked primary AML (n=3 biological replicates). **B)** Barplot showing the fraction of genes bound at the CDS and 3’UTR in the primary AML heat shock condition associated with RNA decay or stabilization from primary AML G3BP1 KD RNA-seq analysis. **C-E)** Average 3’UTR length **C)**, GC content **D)**, and predicted structure **E)** of eCLIP-identified G3BP1-bound transcripts in untreated (control) versus heat shocked primary AML cells. **F)** Analysis of the average 3’UTR length (left), GC content (middle) and structure (right) of eCLIP identified G3BP1 bound transcripts in untreated (Control) versus heat shocked (HS) primary AML cells (N = 3) upregulated or downregulated upon G3BP1 knockdown in primary AML. Significant RNA interactors were selected at P-max <0.05. \*\*\*\**P* <.0001, \*\*\**P* <.001, \*\**P* <.01, \**P* <.05. n.s. not significant.

**Supplementary Figure 8.**
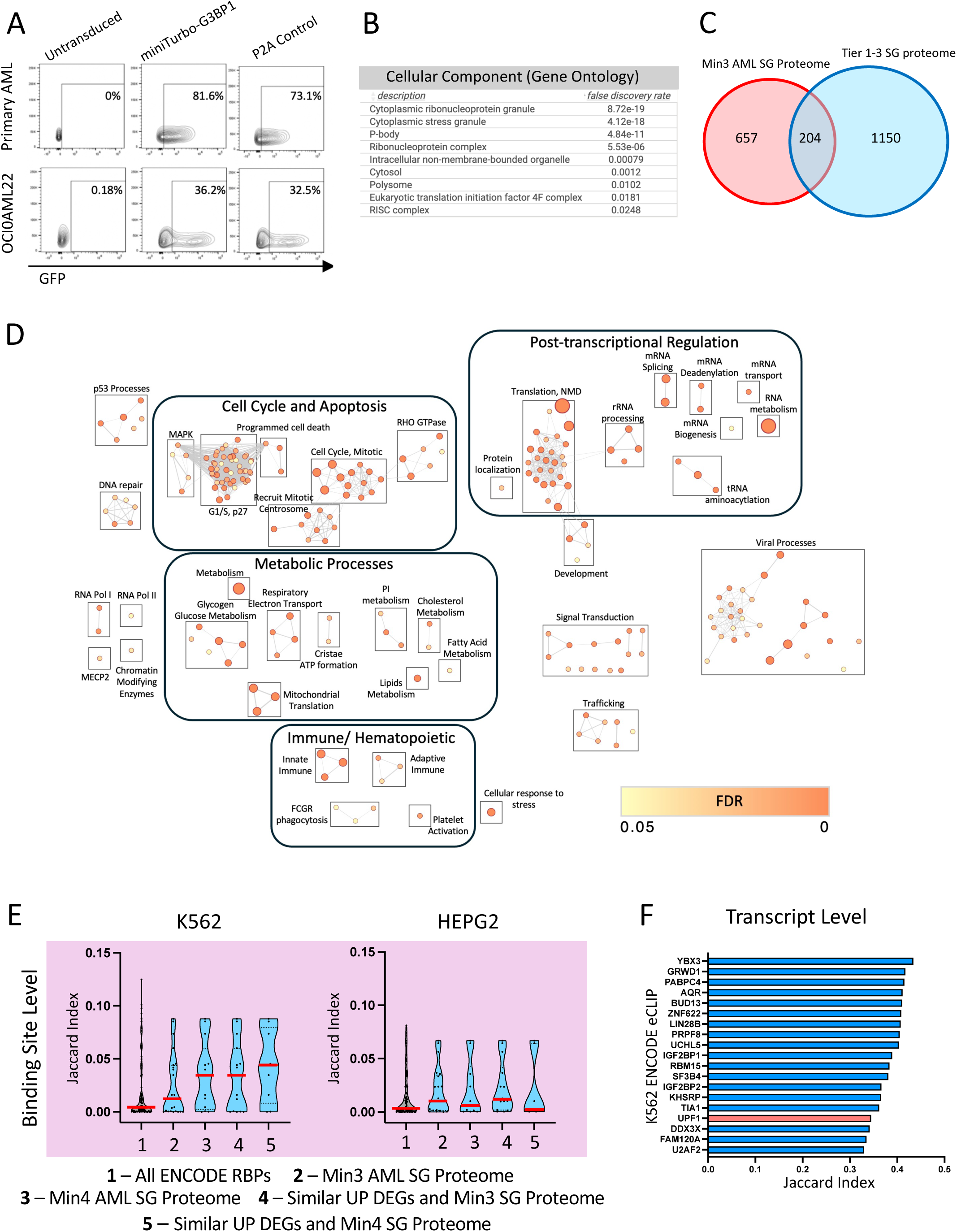
A) Representative flow cytometry plots of miniTurbo overexpression in primary AML and OCI- AML22 cells. **B)** Significant GO Cellular Component terms overrepresented in the core Min6 AML G3BP1 SG proteome determined in STRING. **C)** Venn diagram depicting the overlap in genes identified in the Min3 AML SG proteome versus the Tier 1-3 SG proteome. **D)** Enrichment map of significantly overrepresented Reactome pathways in Min 3 AML SG proteome. Node sizes are proportional to relative normalized enrichment score (NES). **E)** Jaccard index scores comparing heat shocked G3BP1 eCLIP transcript binding site profiles to ENCODE^44^ K562 or HEPG2 CLIP binding profiles in the following categories: 1) All ENCODE RBPs 2) Min 3 AML SG proteome 3) Min 4 AML SG proteome 4) Significant enrichment (GSEA) of upregulated sg/shRBP DEGs within our shG3BP1 upregulated DEGs and Min 3 SG Proteome 5) Significant enrichment (GSEA) of upregulated sg/shRBP DEGs within our shG3BP1 upregulated DEGs and Min4 AML SG Proteome. **F)** Top 15 K562 ENCODE RBPs holding a similar CLIP profile to THP-1 heat shocked cells.

**Supplementary Figure 9.**
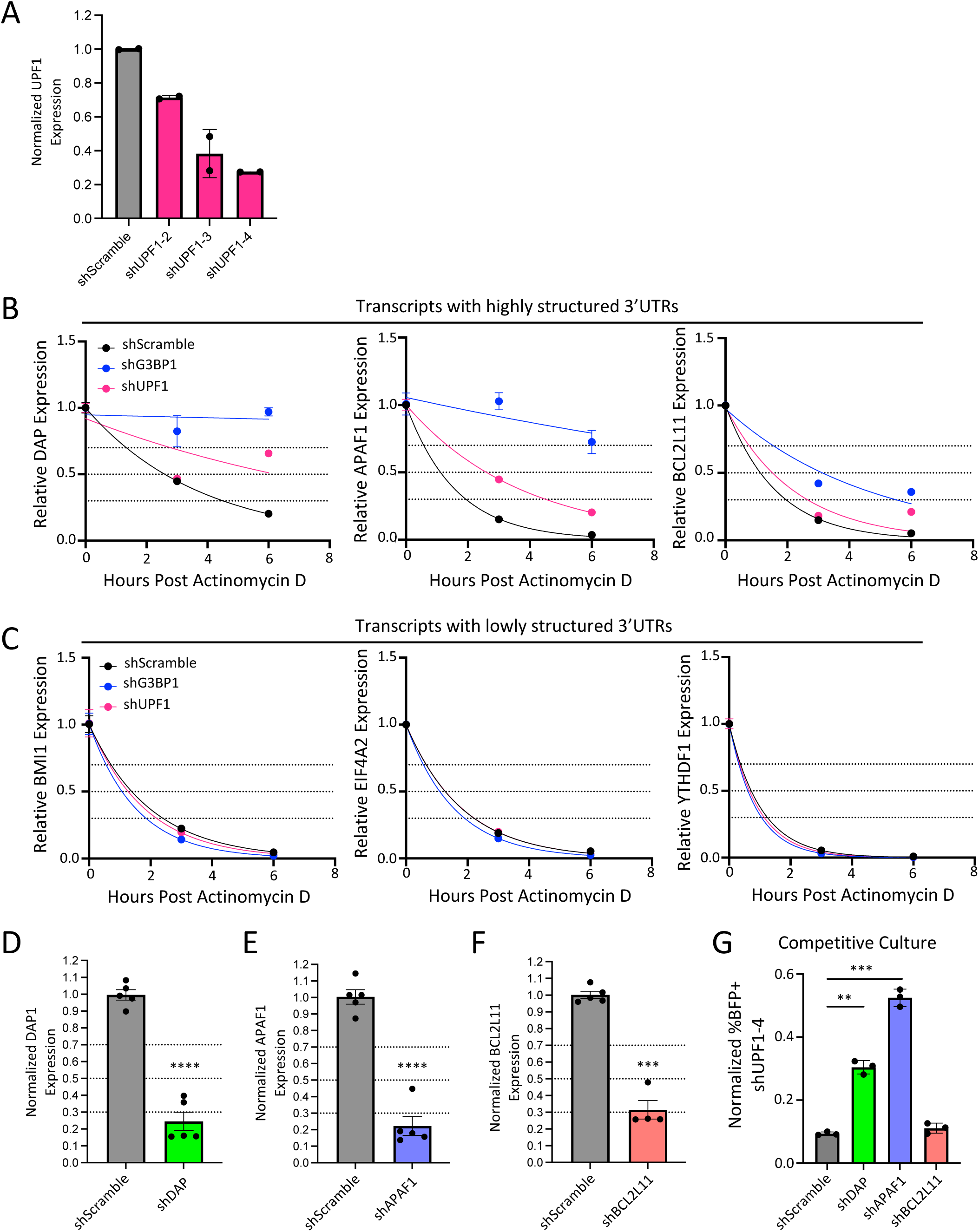
A) qPCR validation of UPF1 knockdown in LentiX cells transfected with shScramble (Control) shUPF1-2, shUPF1-3, or shUPF1-4. Results are normalized to shScramble and percent transfection. **B)** Highly structured 3’UTR APAF1 (left), DAP (middle), and BCL2L11 (right) RNA decay plots in shScramble, shG3BP1-4, or shUPF1-2 knockdown THP-1 cells 0, 3, and 6 hours post Actinomycin D treatment. **C)** Low 3’UTR structured BMI1 (left), EIF4A2 (middle), and YTHDF1 (right) RNA decay plots in shScramble, shG3BP1-4, or shUPF1-2 knockdown THP-1 cells 0, 3, and 6 hours post Actinomycin D treatment. **D-F)** qPCR validation of target knockdown from shDAP **D)**, shAPAF1 **E)** and shBCL2L11 **F)** PLKO.1 lentivirus. **G)** THP-1 shUPF1-4 Day 3 post double transduction competitive rescue assays with additional DAP, BCL2L11, APAF1, or shScramble knockdown. ****P <.0001, ***P <.001, **P <.01, *P <.05. n.s., not significant.

